# A Comprehensive Catalog of Telomere Content Variation Across Human Populations

**DOI:** 10.1101/2025.11.03.686324

**Authors:** Priyanshi Shah, Arun Sethuraman

**Affiliations:** Department of Biology, San Diego State University, San Diego, California, 92182, United States of America

## Abstract

Quantifying telomere content across global human populations is critical to understanding the evolutionary biology of senescence and has implications for the field of biogerontology. Here, we analyzed 26 global human populations from Phase 3 of the 1000 Genomes Project and observed that differences in telomere content vary significantly across different human populations. Most notably, African-ancestry populations in the Americas: ACB (African Caribbean in Barbados) and ASW ((African Ancestry in Southwest USA) showed significantly different telomere content compared to both continental African populations and populations from other ancestries (p < 10^-5^) However, we did not observe significant differences (p > 0.05) in telomere content between sexes across and within different populations. We also report that out of the five superpopulations (separated by continental ancestry), humans of American ancestry show significant difference in telomere content with all other superpopulations (p < 0.001 for all comparisons), whereas humans of East Asian and South Asian ancestry show no significant differences with other superpopulations (p > 0.05). Our analysis suggests that the extent of telomere content variation is likely influenced by complex interactions between genomic and environmental factors. The pipeline to catalog telomere content variation is publicly available via: https://github.com/paribytes/TeloTales.

## INTRODUCTION

The natural, time-dependent deterioration of physiological organ function that ultimately results in death is called aging (Aunan et al., 2016). According to (Franceschi et al., 2018), most pathological conditions that shorten life expectancy and cause chronic illnesses that impact the elderly are primarily caused by aging. These conditions include immune-senescence, cardio-metabolic disorders, osteoporosis, sarcopenia, arthritis, cataracts, neurodegenerative diseases, and many cancers. There is no singular explanation as to why we age. However, the characteristics of aging can be categorized into three main groups (Aunan et al., 2016). The first group, known as primary hallmarks, includes factors that damage cellular functions, such as genomic instability, telomere shortening, changes in gene regulation, and loss of proteostasis (Aunan et al., 2016). Subsequently, the body responds to this damage with antagonistic processes, including deregulated nutrient sensing, altered mitochondrial function, and cellular senescence (Aunan et al., 2016). Lastly, integrative hallmarks, such as stem cell depletion and changes in intercellular communication, may further contribute to the clinical effects of aging, such as reduced reserve capacity, organ decline, and decreased function (Aunan et al., 2016). Many of the suggested characteristics are like those found in other diseases like cancer (Aunan et al., 2016).

Telomeres in vertebrates consist of tandem hexameric sequence repeats (TTAGGG)n located at the ends of linear chromosomes (Victorelli & Passos, 2017). These repetitive nucleotide sequences safeguard chromosome ends and uphold genomic stability (Aunan et al., 2016). At a cellular level, telomere dysfunction and genomic instability seem to play a crucial role in the aging process (Xie et al., 2015). Telomere shortening is commonly linked to age-related illnesses and premature aging conditions (Aunan et al., 2016).

Most human somatic cells do not produce telomerase, an enzyme that can lengthen the telomeres (Aunan et al., 2016). Because human DNA polymerase cannot completely replicate the telomere ends of DNA, the DNA strands progressively shorten with each cell division (Aunan et al., 2016). During each division of somatic cells, the telomeres become shorter by 50–200 base pairs (bp) due to incomplete synthesis of the lagging strand during DNA replication (Srinivas et al., 2020). This is caused by the inability of DNA polymerase to fully replicate the 3′end of the DNA strand, a phenomenon known as "the end-replication problem" (Olovnikov, 1973; Watson, 1972). This can eventually affect the coding regions of the DNA (Aunan et al., 2016). As a solution to the “end-replication problem”, cells undergo replicative senescence after a specific number of divisions and as the telomeres shorten, a phenomenon referred to as the Hayflick limit (Aunan et al., 2016). Additionally, the G-rich telomere repeat sequence is highly susceptible to oxidative damage, which can directly damage telomeres, leading to cell senescence (Oikawa & Kawanishi, 1999).

In individuals over 60 years old, having shorter telomeres has been linked to earlier mortality due to age-related illnesses (Aunan et al., 2016). While increased expression of telomerase can slow down aging, it can also lead to tumorigenesis (Aunan et al., 2016). Research involving families with long-living members discovered that telomere length decreases with age in men, indicating a strong hereditary component to telomere length (Aunan et al., 2016). Additionally, telomere shortening has been linked to several age-related diseases, including osteoarthritis, atherosclerosis, coronary heart disease, and atrial fibrillation (Aunan et al., 2016).

Here, we catalog 1000 Genomes Project’s Phase 3 data using telomere content collated from low-coverage whole genomes (mean depth 7.4x) to answer four main research questions: (1) are there significant differences in telomere content across diverse human populations?, (2) does telomere content vary among different continental superpopulations?, (3) does telomere content vary between self-reported sexes within and across populations?, (4) and how does telomere content vary autosome-wise in each population?

## METHODS

### Estimating telomere content

Molecular methods to estimate telomere content and length include experimental techniques like Quantitative polymerase chain reaction (q-PCR), Terminal restriction fragment (TRF) length analysis and Flow-fluorescent *in situ* hybridization (flow-FISH). Alternate approaches utilize computational tools to analyze large-scale, high-coverage whole-genome sequencing (WGS) data. These latter WGS-based techniques provide a robust and accurate alternative to experimental techniques. Recently developed tools to estimate telomere content and length include Motif_counter (Sun et al., 2023), TelSeq (Ding et al., 2014), Computel (Nersisyan & Arakelyan, 2015), qmotif (Holmes et al., 2022), and Telomerecat (Farmery et al., 2018) (Lee et al., 2017). In this study, we use qmotif v1.0 to determine telomere content as an estimate of telomere length directly from WGS data of diverse human populations. qmotif v1.0 is a fast, multi-threaded, and user-friendly tool for quantifying telomere content from WGS data (Holmes et al., 2022). qmotif has also previously been benchmarked to outperform other telomere content and length quantification tools, with a strong correlation with qPCR-based estimates of telomere length. qmotif was also determined to be the fastest amongst tools compared due to its efficient utilization of regular expression based string-matching (Holmes et al., 2022).

To determine the telomere content of each individual, we utilized individual BAM files (Binary Alignment Map) and their corresponding binary index files (BAI) from NCBI’s 1000 Genomes Dataset website (https://www.ncbi.nlm.nih.gov/projects/faspftp/1000genomes/). The BAM files from Phase 3 were mapped to GRCh37/hg19 reference genome build. Files were downloaded for each sample and used as input for the qmotif v1.0 tool, with the provision of a qmotif configuration file for hg19 telomere quantification. This file follows the standard INI-file format and has three sections: PARAMS, INCLUDES, and EXCLUDES. The PARAMS section outlines the search criteria and customizable parameters (e.g., window size, cutoff size). The INCLUDES section contains a list of genomic regions intended for analysis (e.g., telomeric regions) (see appendix Table S1). The EXCLUDES section contains a list of genomic regions that are to be excluded from the analysis (e.g., centromeres or other genomic regions with extensive repeat content).

Following our pipeline, all statistical analyses described below were conducted using R (R Core Team, 2024) in RStudio (RStudio Core Team, 2024) using the ggplot2 v3.5.1 (Wickham, 2016), dplyr v1.1.4 (Wickham et al., 2023), tidyverse v2.0.0 (Wickham et al., 2019), colorspace v2.1.1 (Zeileis et al., 2009), car v3.1.3 (Fox et al., 2019) and rstatix v0.7.2 (Kassambara, 2023) packages.

### Statistical analyses of variation in telomere content

Due to the large sample-size of human genomes analyzed, we first used a Shapiro-Wilk test followed by a one-sample Kolmogorov-Smirnov test to check for normality in telomere content across populations. For both the normality tests, the hypotheses are as follows: The null hypothesis (H0) proposes that the sample data, here telomere content, follows a normal distribution, whereas the alternative hypothesis (H1) is that telomere content does not follow the normal distribution. Then we performed log-transformation on the telomere content data to address the positive skewness in the data. Since the one-way ANOVA test assumes homogeneity of variance, we used Levene’s test to verify equality of variances in telomere content across the populations. Since the homogeneity of variance assumption was violated, instead of one-way ANOVA, we used a one-way Welch ANOVA test which is a robust alternative to the standard ANOVA that does not assume equal variances across groups. The hypotheses for Welch’s ANOVA test are as follows: The null hypothesis (H0) is that the population means are all equal, whereas the alternative hypothesis (H1) is that the population means are not equal. Then as a post-hoc test for Welch’s ANOVA to identify which specific populations differ from each other, we performed a Games-Howell test, which is designed for situations with unequal variances. The hypotheses for Games-Howell test are as follows: The null hypothesis (H0) proposes that there is no difference in the mean values of telomere content between the two groups being compared. The alternative hypothesis (H1) states that there is a difference in the mean values of telomere content data between the two groups being compared.

We then repeated the statistical tests described above to test for significant differences in telomere content after grouping individuals into their continental ancestries. Thereon, we applied Welch’s ANOVA test to compare telomere content variation across multiple groups, followed by a Games-Howell test to determine statistically significant variation across our designated superpopulations.

In order to test for differences in telomere content across sexes, after grouping telomere content variation across self-reported sexes, we performed a Welch two sample t-test to test for differences in telomere content between self-reported sexes across all the populations. The hypotheses for the Welch two sample t-test are as follows: The null hypothesis (H0) proposes that there is no difference in the means of telomere content between males and females. The alternative hypothesis (H1) states that there is a difference in the means of telomere data between males and females and the mean of telomere content is not equal between sexes.

We also used a Welch two sample t-test was to test for significant differences in telomere content between self-reported sexes within each population. P-value correction for multiple testing was then performed using the conservative Bonferroni method.

## RESULTS

All available 2353 WGS BAM files from Phase 3 of the 1000 Genomes Project were analyzed using the pipeline described above (Fig. 2). Out of the 2353 samples, 1196 samples belonged to self-reported females and 1157 belonged to self-reported males across 26 global human populations.

**Fig 1.**
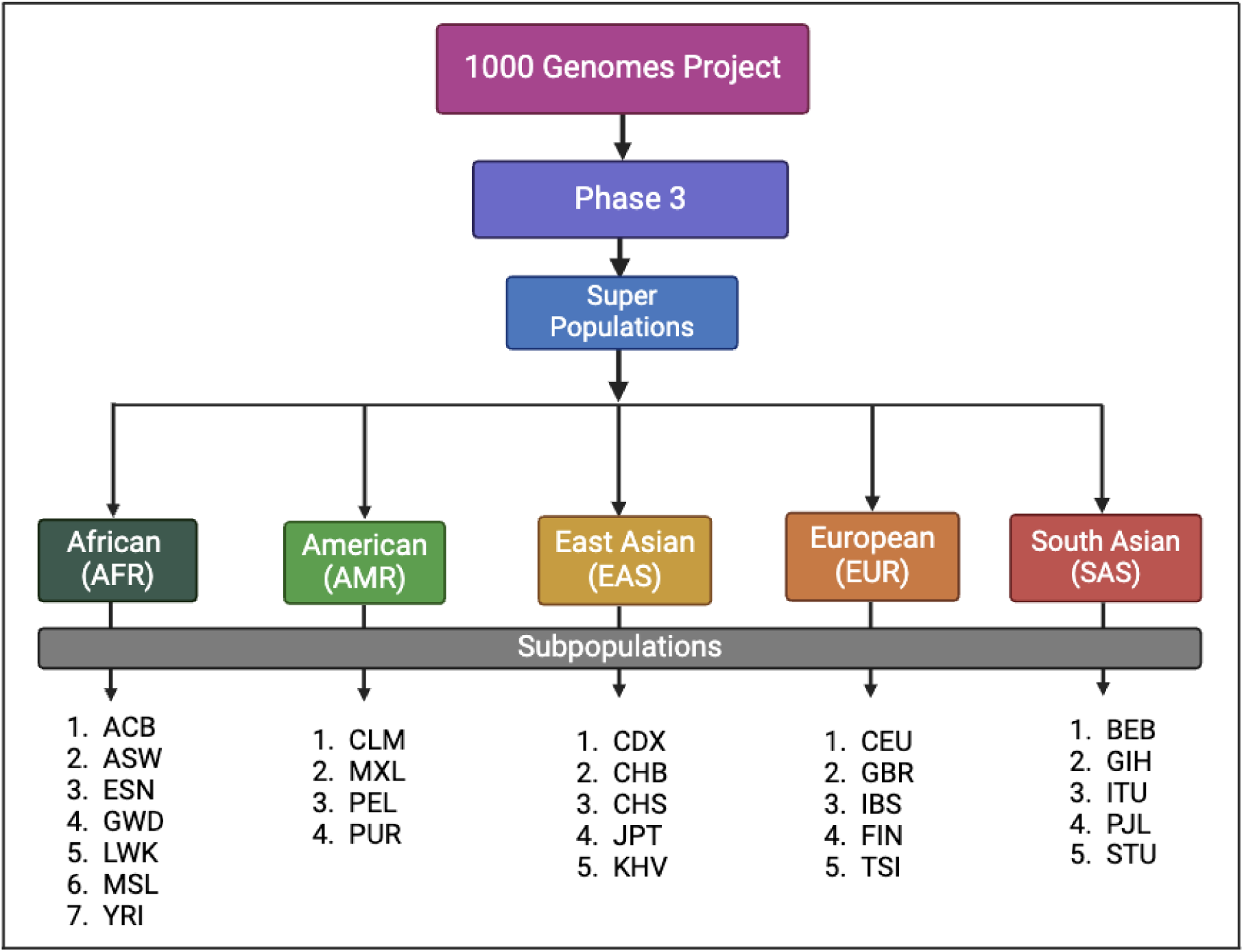
**Hierarchical scheme of superpopulations and their corresponding subpopulations in Phase 3 of the 1000 Genomes Project.** The 1000 Genomes Project has 26 global populations, organized into five superpopulations based on continental ancestry: African (AFR, n=7 populations), American (AMR, n=4 populations), East Asian (EAS, n=5 populations), European (EUR, n=5 populations), and South Asian (SAS, n=5 populations). Population codes are defined in Table 2. This hierarchical structure was used for both superpopulation-level and population-level analyses of telomere content variation.

**Fig 2.**
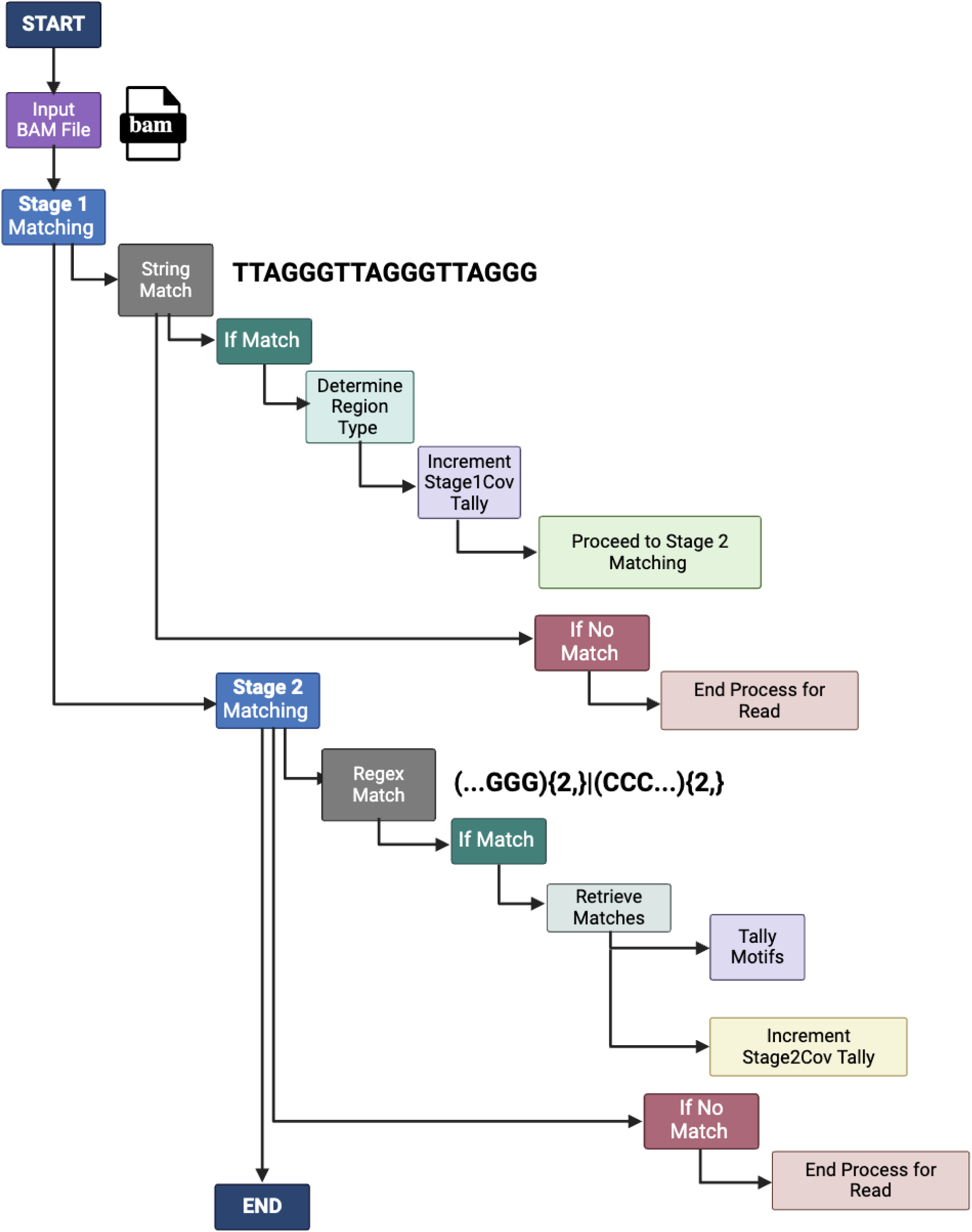
Workflow for our telomere content variation pipeline, that incorporates qmotif to estimate variation from across Phase 3 of the 1000 Genomes Project. Schematic representation of the qmotif-based pipeline used to estimate telomere content variation across samples from the 1000 Genomes Project. Our workflow involves extracting and quantifying telomeric reads from whole-genome sequencing data using qmotif. qmotif operates through a two-stage matching system: in Stage 1, a simple string match is used to identify canonical telomeric repeats, while in Stage 2, a more complex regular expression is applied to detect variant telomeric sequences. At the end of Stage 2, a tally of all identified motifs is collated, and the final number is recorded. [Figure created with BioRender.com].

**Fig 3.**
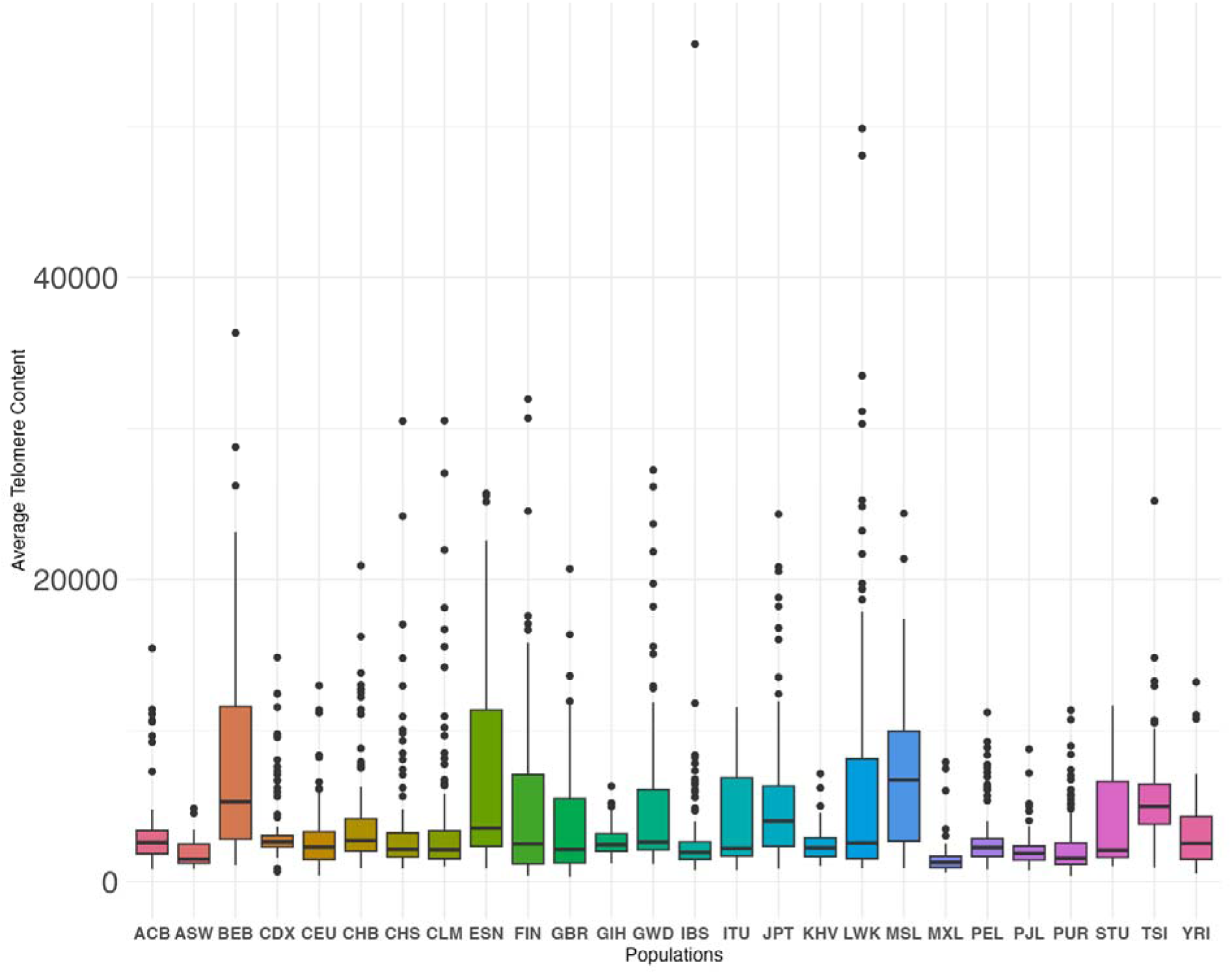
Telomere content distribution across 26 global populations. Box-and-whisker plot showing the distribution of average telomere content across 26 populations from Phase 3 of the 1000 Genomes Project (population IDs described in Table 2). Boxes show interquartile range (IQR), lines show medians, whiskers extend to 1.5×IQR, points are outliers. One-way ANOVA revealed significant variation across populations (*F [25, 819.09] = 22.629, p < 2.2e-16),* with pairwise comparisons identifying significant differences between multiple population pairs after correction for multiple testing (see appendix Table S2 for all comparisons).

**Fig 4.**
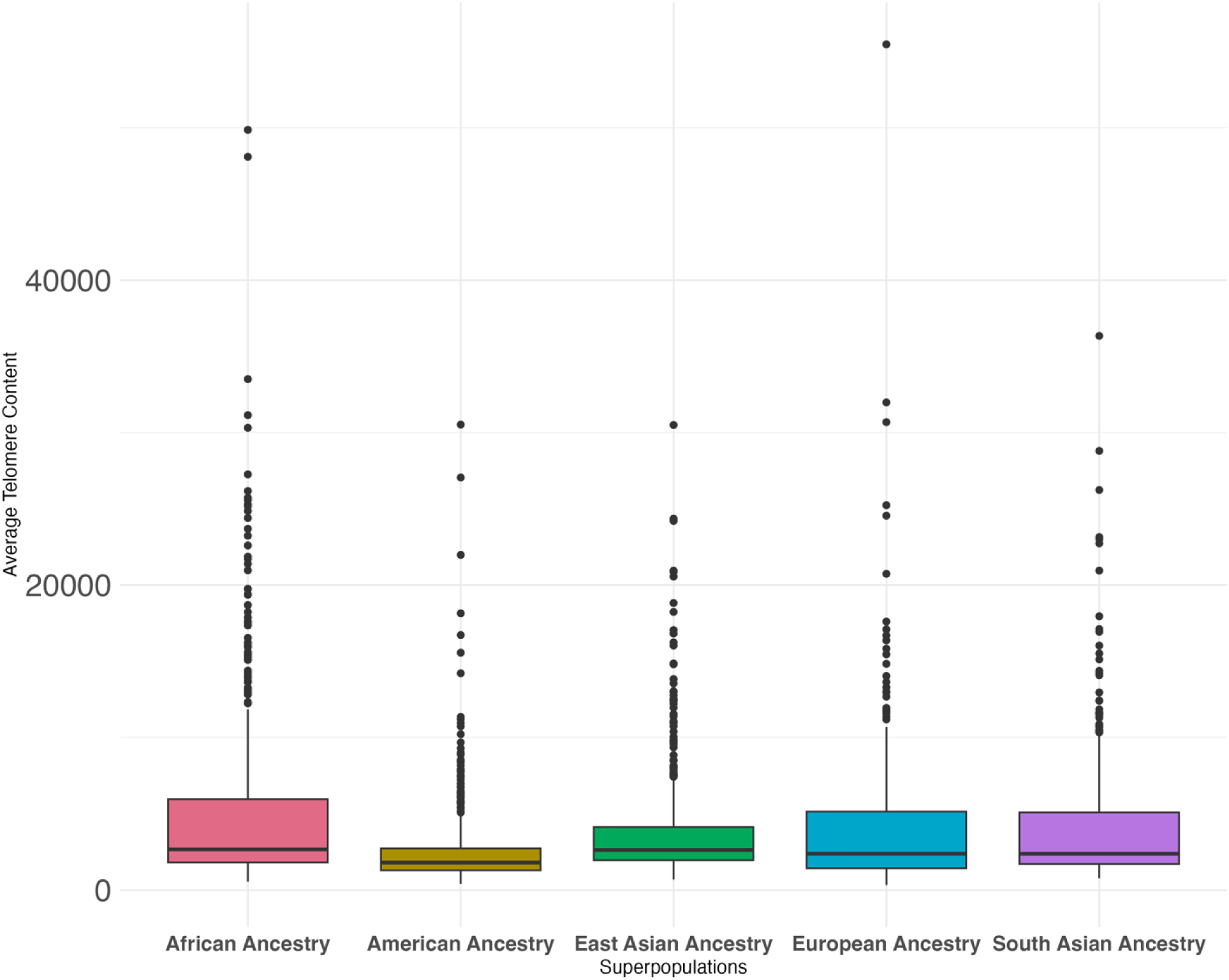
Telomere content distribution across 5 superpopulations. Box-and-whisker plot of average telomere content from Phase 3 of the 1000 Genomes Project across five superpopulations. Boxes show IQR, lines show medians, whiskers extend to 1.5×IQR, points are outliers. American ancestry populations show significant difference when compared to all other superpopulations (p < 0.001 for all comparisons). Superpopulation descriptions are in Table 1.

**Fig 5.**
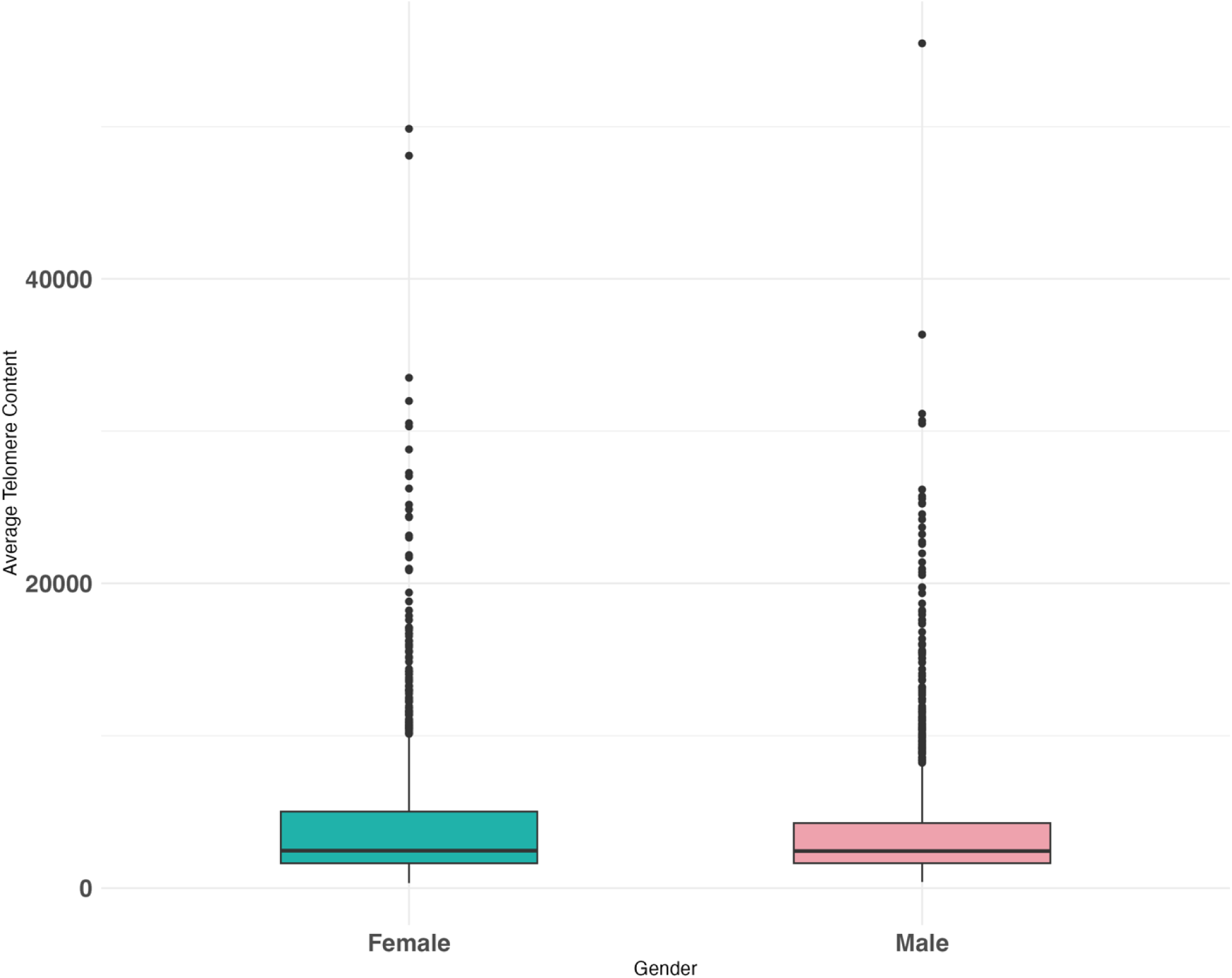
Telomere content distribution by sex across all 26 populations. Box-and-whisker plot comparing average telomere content between females (n = 1196) and males (n = 1157) from Phase 3 of the 1000 Genomes Project. Boxes show IQR, lines show medians, whiskers extend to 1.5×IQR, points are outliers. No significant differences were observed between sexes either overall (p = 0.2744) or within individual populations (all p > 0.05), indicating that sex does not significantly influence telomere content in this dataset.

**Fig 6.**
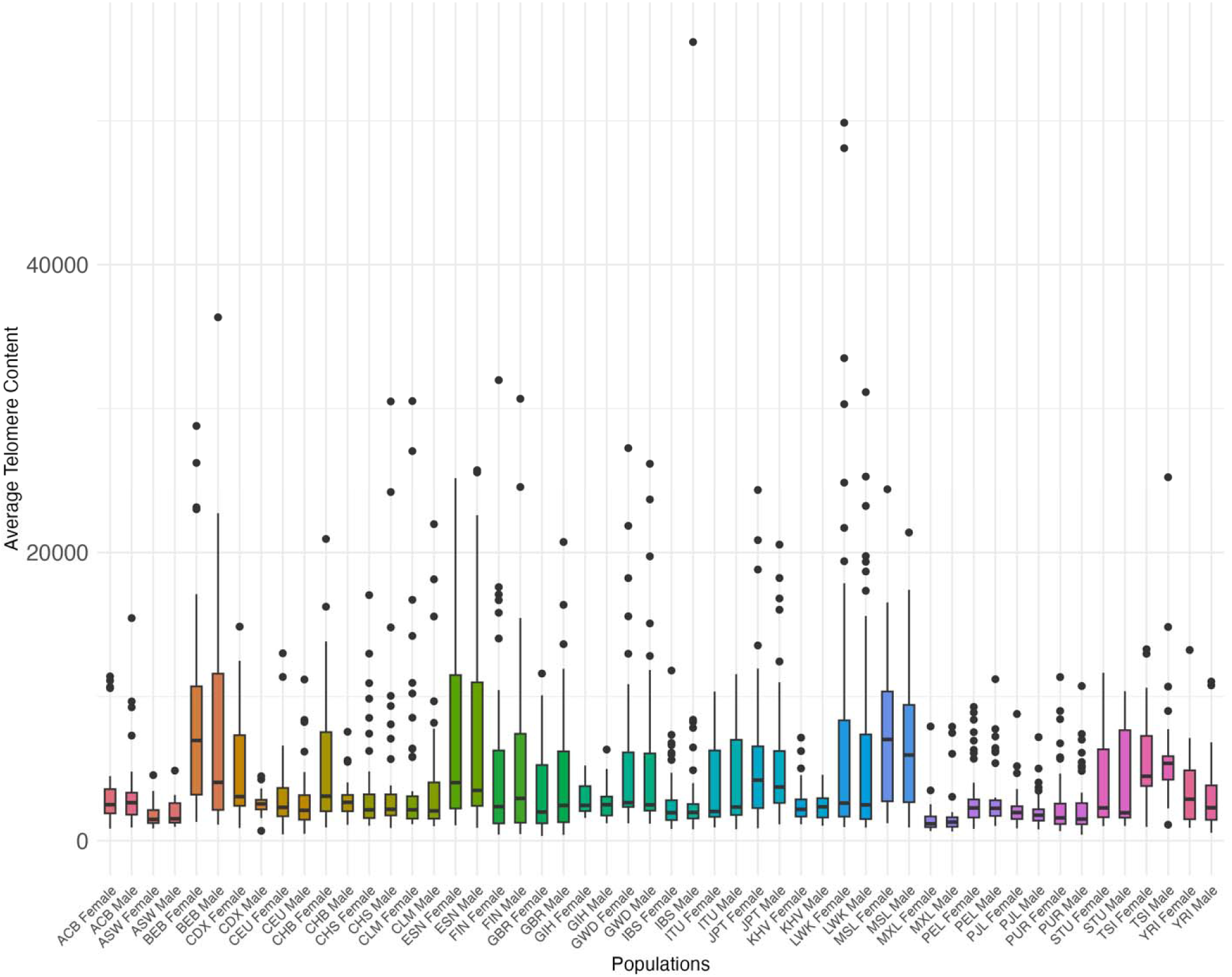
Telomere content distribution by sex within population for all 26 populations. Box-and-whisker plot of average telomere content in self-reported males and females within population across 26 global populations from Phase 3 of the 1000 Genomes Project. Boxes show IQR, lines show medians, whiskers extend to 1.5×IQR, points are outliers. No significant sex differences were detected within any population (all adjusted p > 0.05) or overall (p = 0.2744). Population codes are in Table 2.

Mean telomere content computed across 26 populations was estimated to be 4146.936 (SD 4702.16); with a median of 2454; range 334-55479. A high standard deviation indicates high variability in telomere content across all the populations. The difference between the mean (4146.936) and median (2454) suggests that the data might be right- skewed, with some very high estimated telomere content values pulling the mean upwards.

The Shapiro-Wilk test of normality in telomere content across populations indicated significant deviation from normality (*W=0.62968, p < 2.2e-16*). The Kolmogorov-Smirnov test of normality also revealed a statistically significant deviation from a normal distribution (*D (25) = 0.24039, p < 2.2e-16*). Then we performed log-transformation on the dataset. The Levene’s test was performed on the log normally distributed dataset which indicated that the assumption of homogeneity of variance is violated in telomere content of populations. A one-way Welch ANOVA was conducted to evaluate the differences in log-transformed telomere content among different populations. The results revealed a statistically significant effect of populations on telomere content, (*F[25, 819.09] = 22.629, p < 2.2e-16)*. We, therefore, reject the null hypothesis that there is no significant difference in telomere content across the populations examined.

Further post-hoc analysis with Games-Howell test showed significant differences in the distributions of telomere content between populations (see appendix Table S2) across the globe.

To determine if the telomere content varied significantly among the five superpopulations, we first performed a one-way Welch ANOVA which indicated significant differences (*p < 0.05*) among the five superpopulations, meaning the distributions of telomere content are not the same across the superpopulations. Thereon, we performed a Games-Howell post-hoc test to determine variation between superpopulations, which indicated statistically significant (*p < 0.05*) differences in telomere content across several superpopulations (see appendix Table S3).

Telomere content variation across individuals with American ancestry shows significant difference with all other superpopulations. Whereas individuals of East Asian ancestry and South Asian ancestry show no significant differences with several other superpopulations. Additional pairwise comparisons - African vs East Asian ancestry, East Asian Ancestry vs European Ancestry, African vs South Asian ancestry, East Asian vs South Asian Ancestry and European vs South Asian Ancestry indicate non-significant differences in telomere content variation (see appendix Table S3)

To determine the statistical significance in telomere content variation between males and females across all populations, we performed a Welch’s t-test. With a *p*-value of 0.2744, we fail to reject the null hypothesis, indicating that there is no statistically significant difference in telomere data between self-reported males and females across all the populations.

To determine the statistical significance in telomere content variation between males and females within the same population, we performed a Welch’s t-test. The results of the analysis indicate that only individuals from the CDX population (Chinese Dai in Xishuangbanna) and the CHB population (Han Chinese in Beijing) demonstrated statistically significant differences in telomere content between genders. The *p*-values for these groups were 0.00019 and 0.0032, respectively indicating a sex-based difference in telomere content distribution within that population. All other populations (e.g., GBR, FIN, PUR, see appendix Table S4), the *p*-values were greater than the significance threshold of 0.05, suggesting that there is no statistically significant difference in telomere content between self-reported males and females within these populations. However, correcting for multiple-testing yielded an adjusted false positive rate cutoff α *< 0.00192* discounted both CDX and CHB populations from any statistically significant differences in sex-based telomere content.

Finally, quantifying variation in average telomere content distributions across autosomes within each population (Figs. 7–32) indicated a higher degree of variation, and higher median telomere content across chromosomes 5 and 12, while content remained consistently lower across all other autosomes.

**Fig 7.**
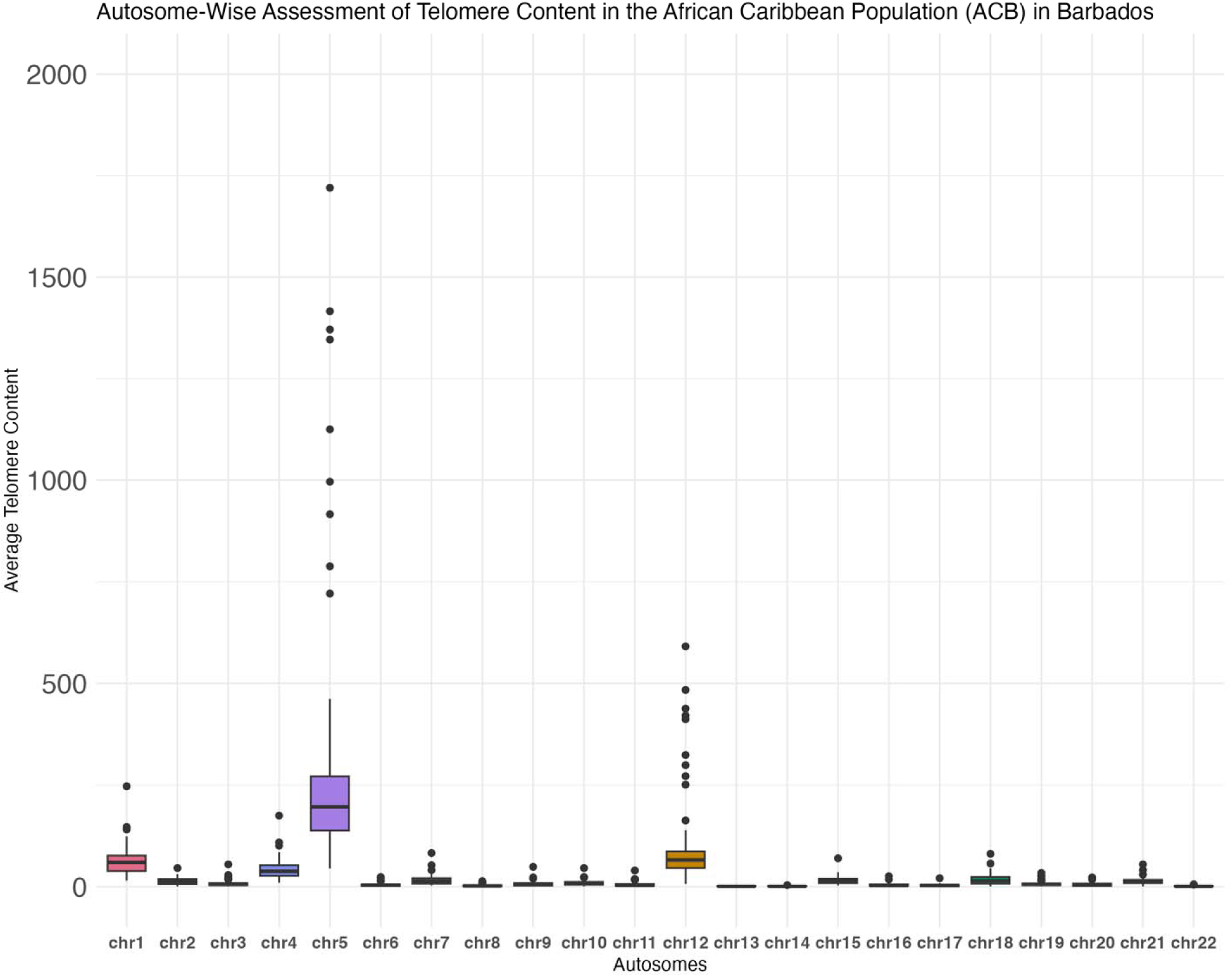
**Autosome-wise telomere content distribution in the African Caribbean population in Barbados (ACB).** Box-and-whisker plot of autosome-wise assessment of average telomere content per autosome in ACB (n = 88).

**Fig 8.**
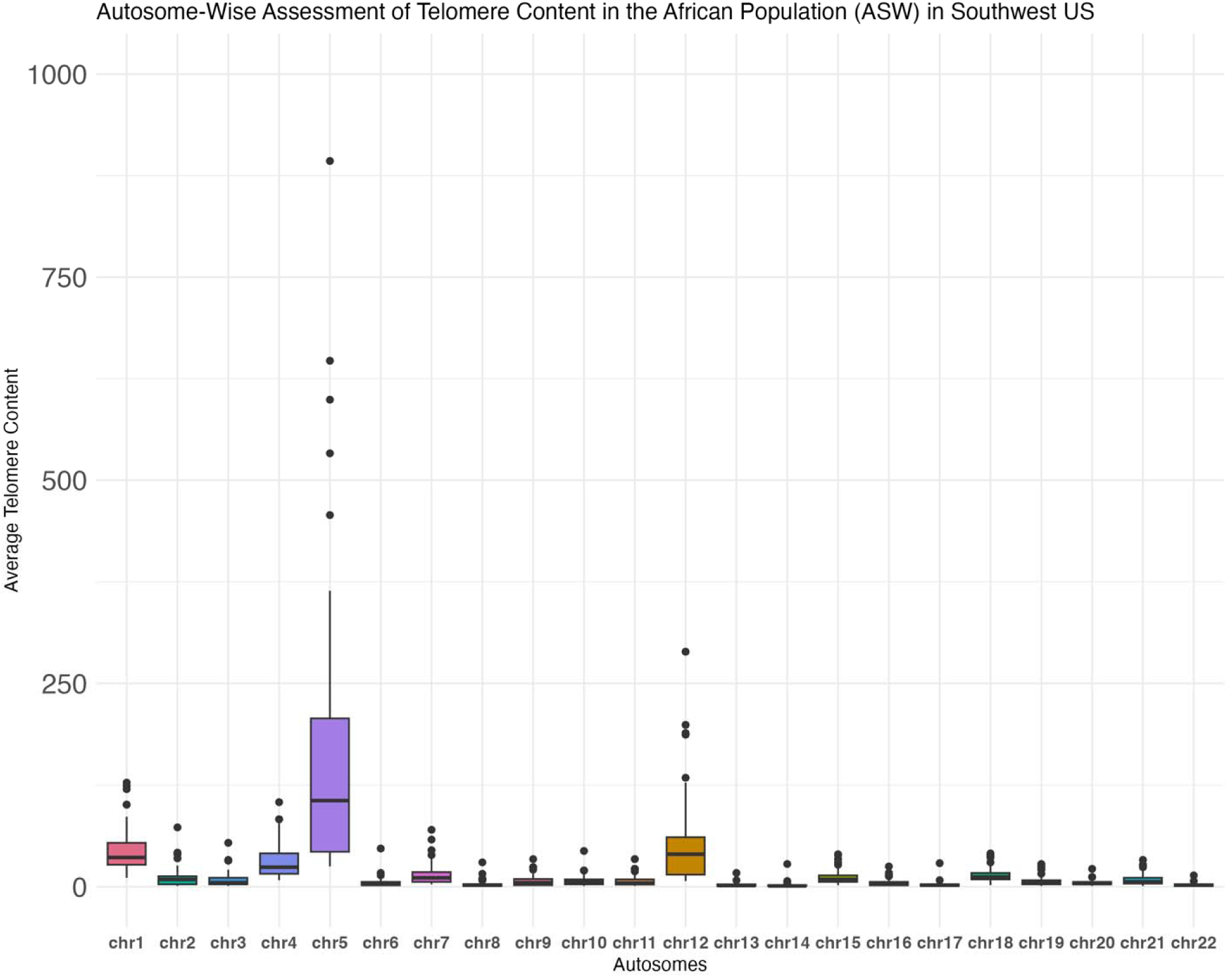
Autosome-wise telomere content distribution in the African population (ASW) in Southwest, USA. Box-and-whisker plot of autosome-wise assessment of average telomere content per autosome in ASW (n = 65).

**Fig 9.**
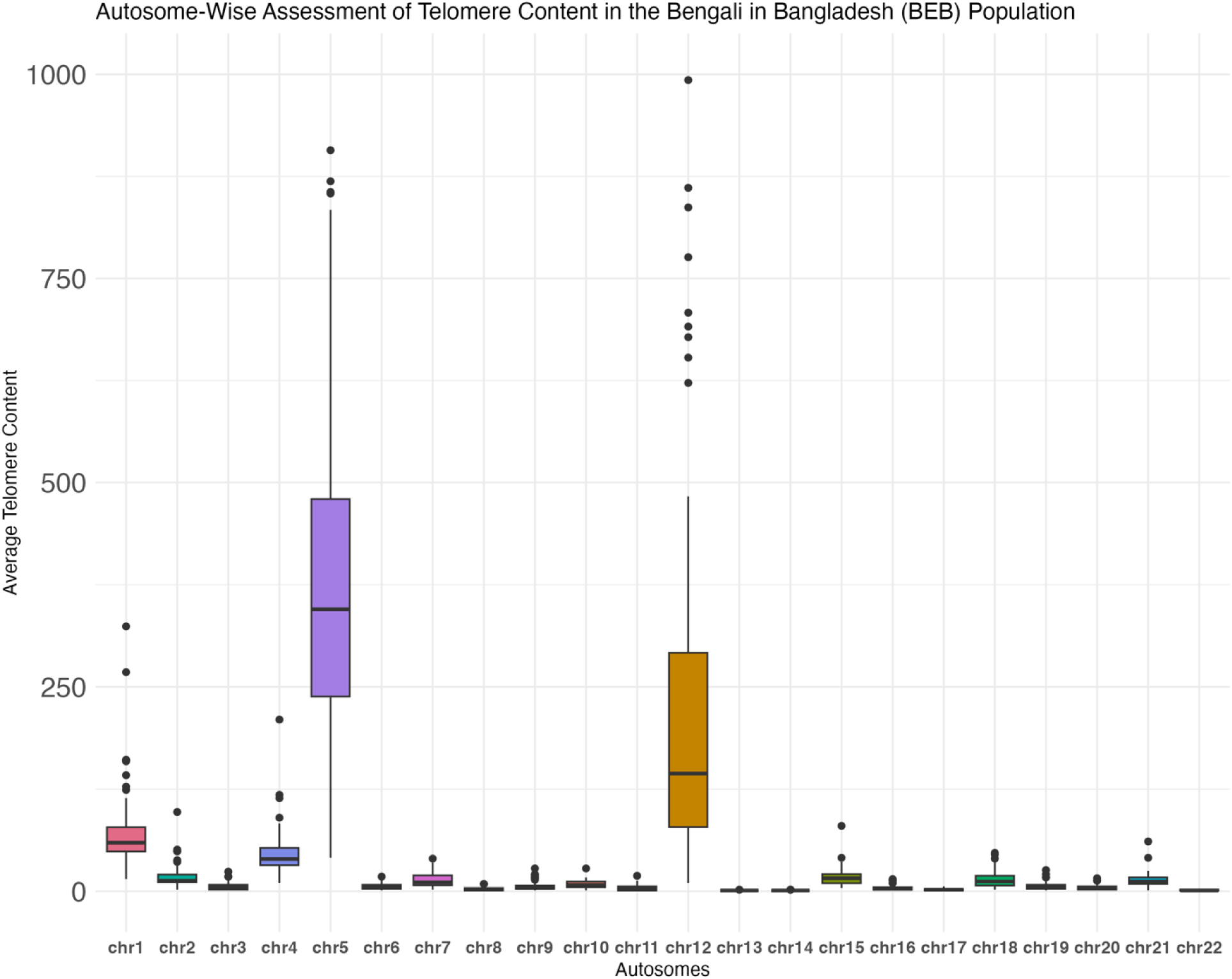
Autosome-wise telomere content distribution in the Bengali population in Bangladesh (BEB). Box-and-whisker plot of autosome-wise assessment of average telomere content per autosome in BEB (n = 84).

**Fig 10.**
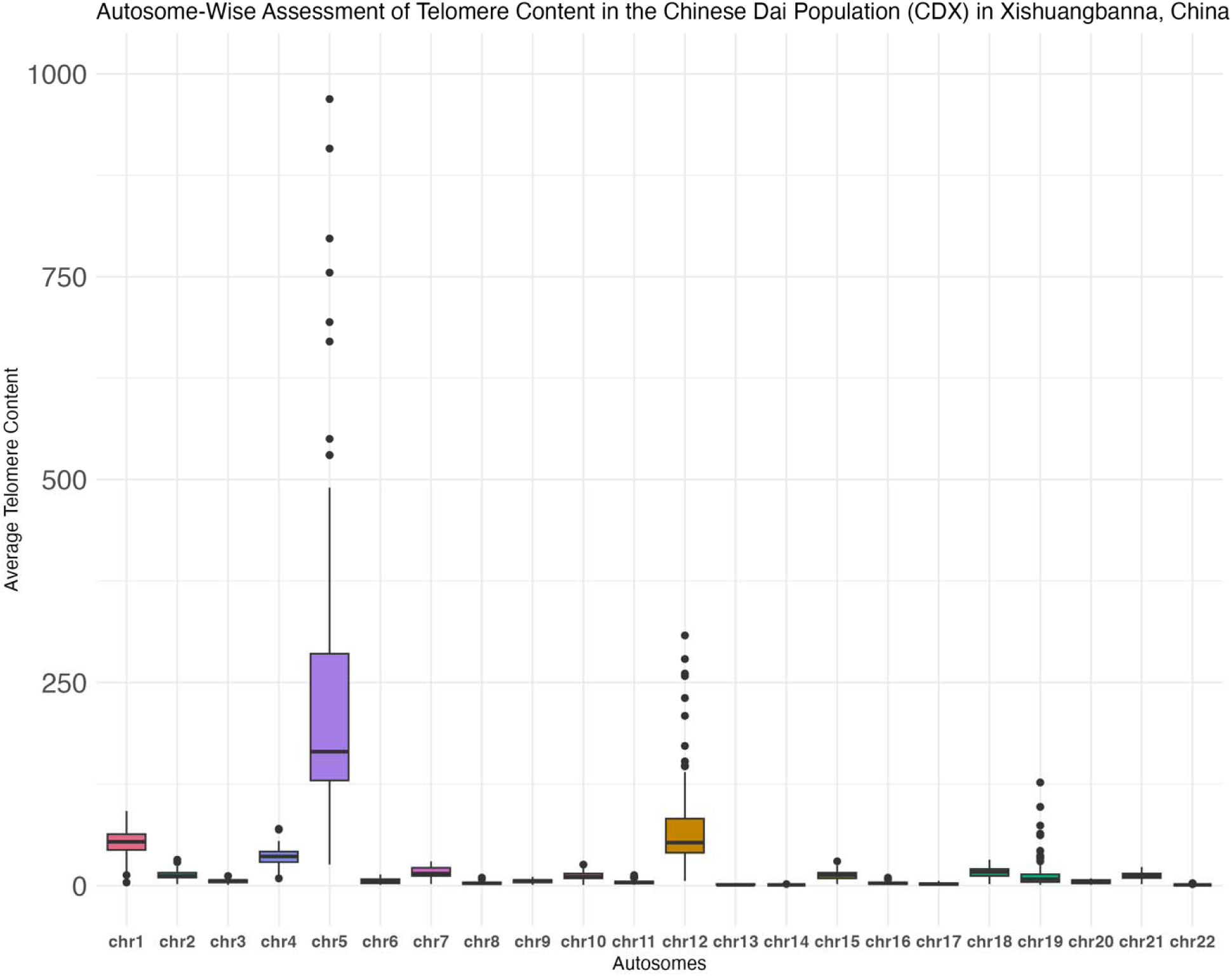
Autosome-wise telomere content distribution in the Chinese Dai population (CDX) in Xishuangbanna, China. Box-and-whisker plot of autosome-wise assessment of average telomere content per autosome in CDX (n = 83).

**Fig 11.**
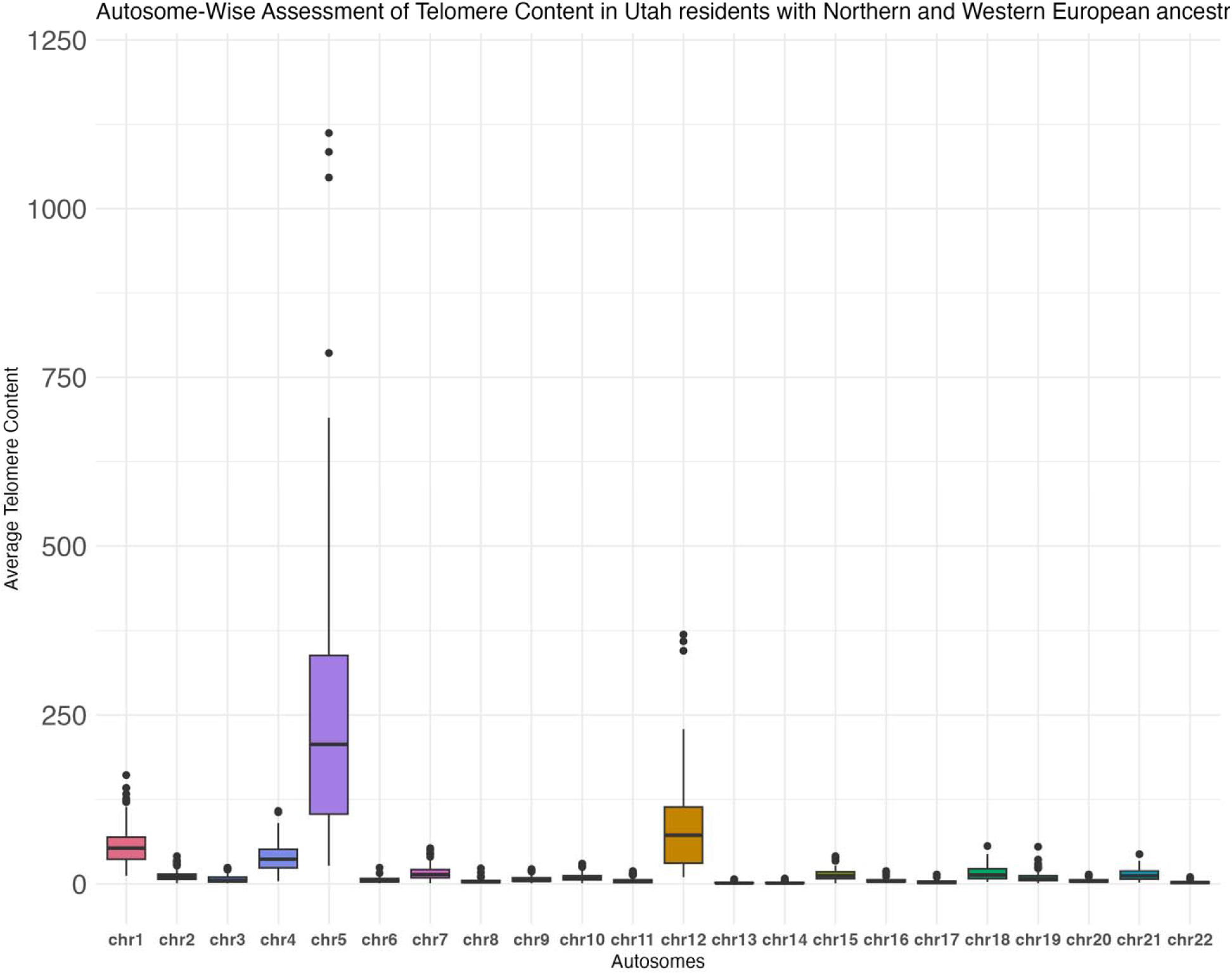
Autosome-wise telomere content distribution in the Utah population with Northern and Western European ancestry (CEU). Box-and-whisker plot of autosome-wise assessment of average telomere content per autosome in CEU (n = 92).

**Fig 12.**
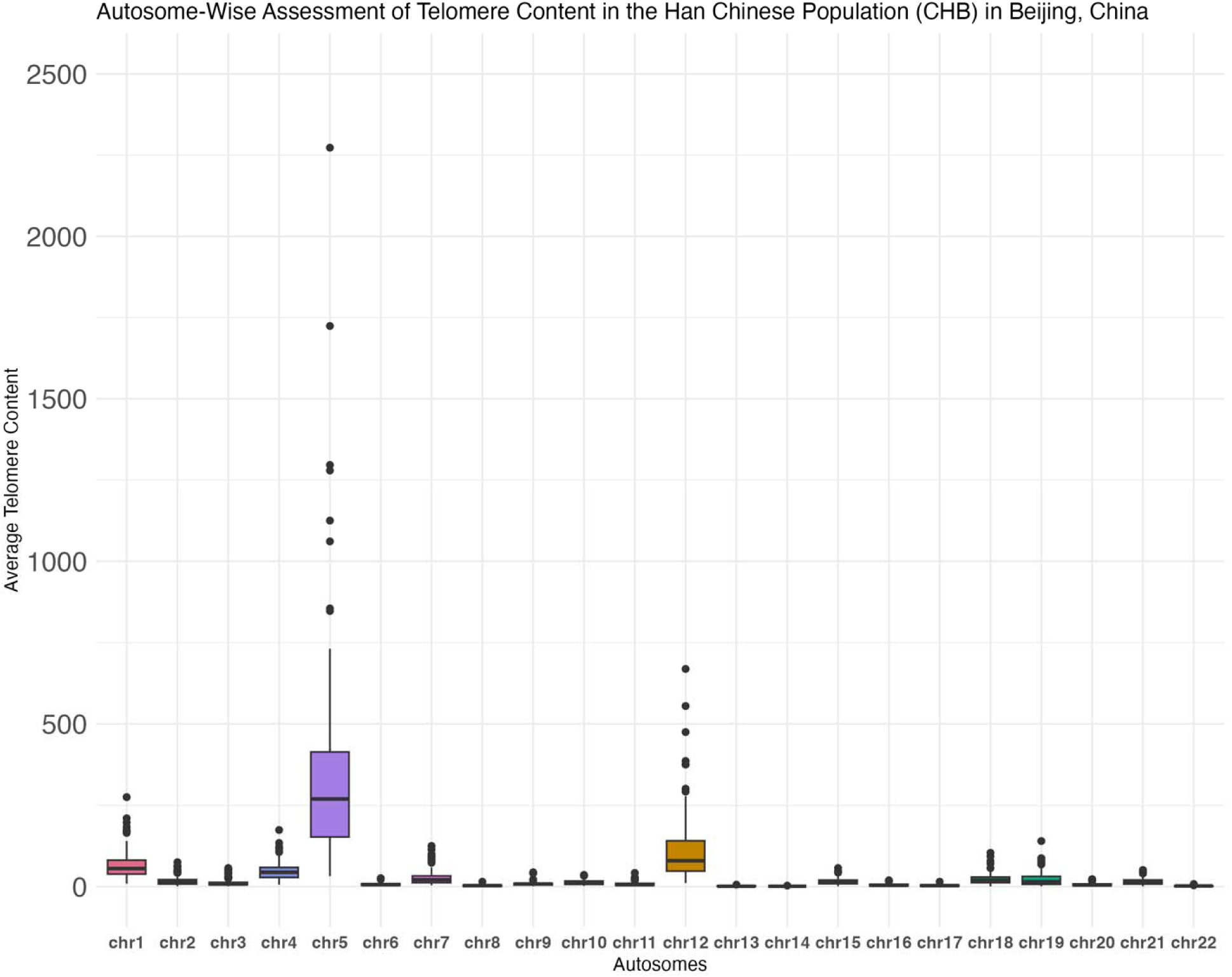
Autosome-wise telomere content distribution in the Han Chinese population (CHB) in Beijing, China. Box-and-whisker plot of autosome-wise assessment of average telomere content per autosome in CHB (n = 101).

**Fig 13.**
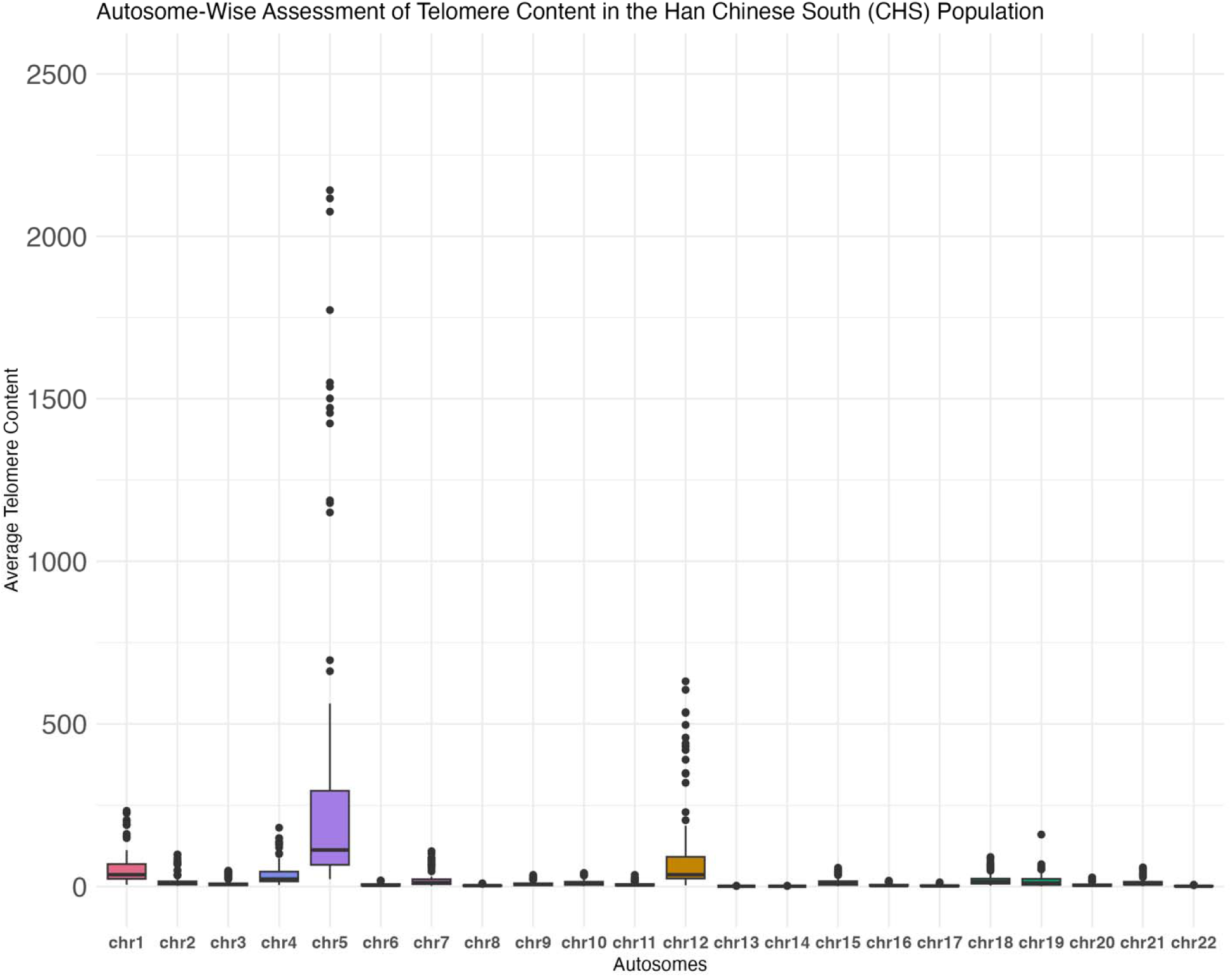
Autosome-wise telomere content distribution in the Han Chinese South (CHS) population. Box-and-whisker plot of autosome-wise assessment of average telomere content per autosome in CHS (n = 104).

**Fig 14.**
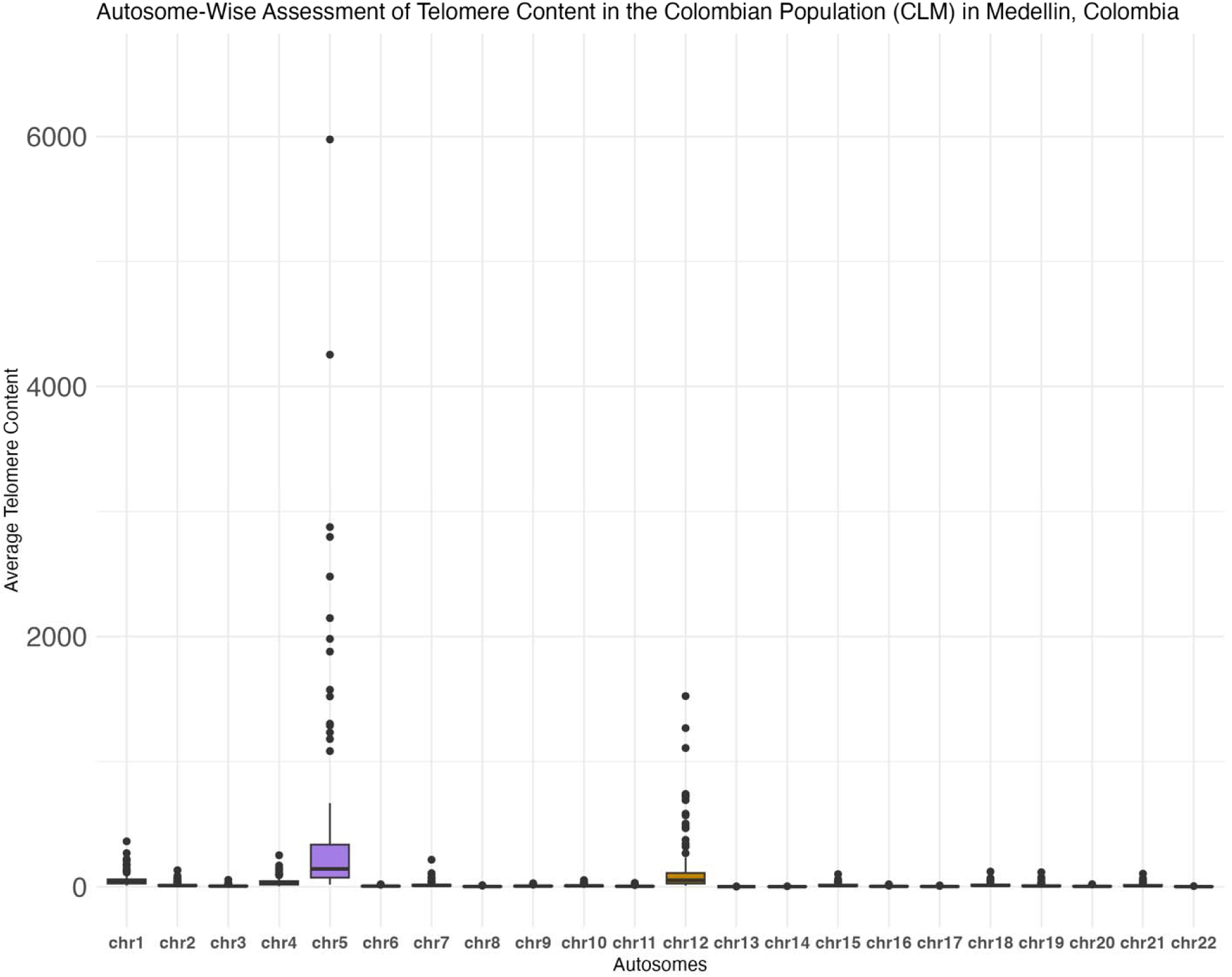
Autosome-wise telomere content distribution in the Colombian population in Mendellin Columbia (CLM). Box-and-whisker plot of autosome-wise assessment of average telomere content per autosome in CLM (n = 94).

**Fig 15.**
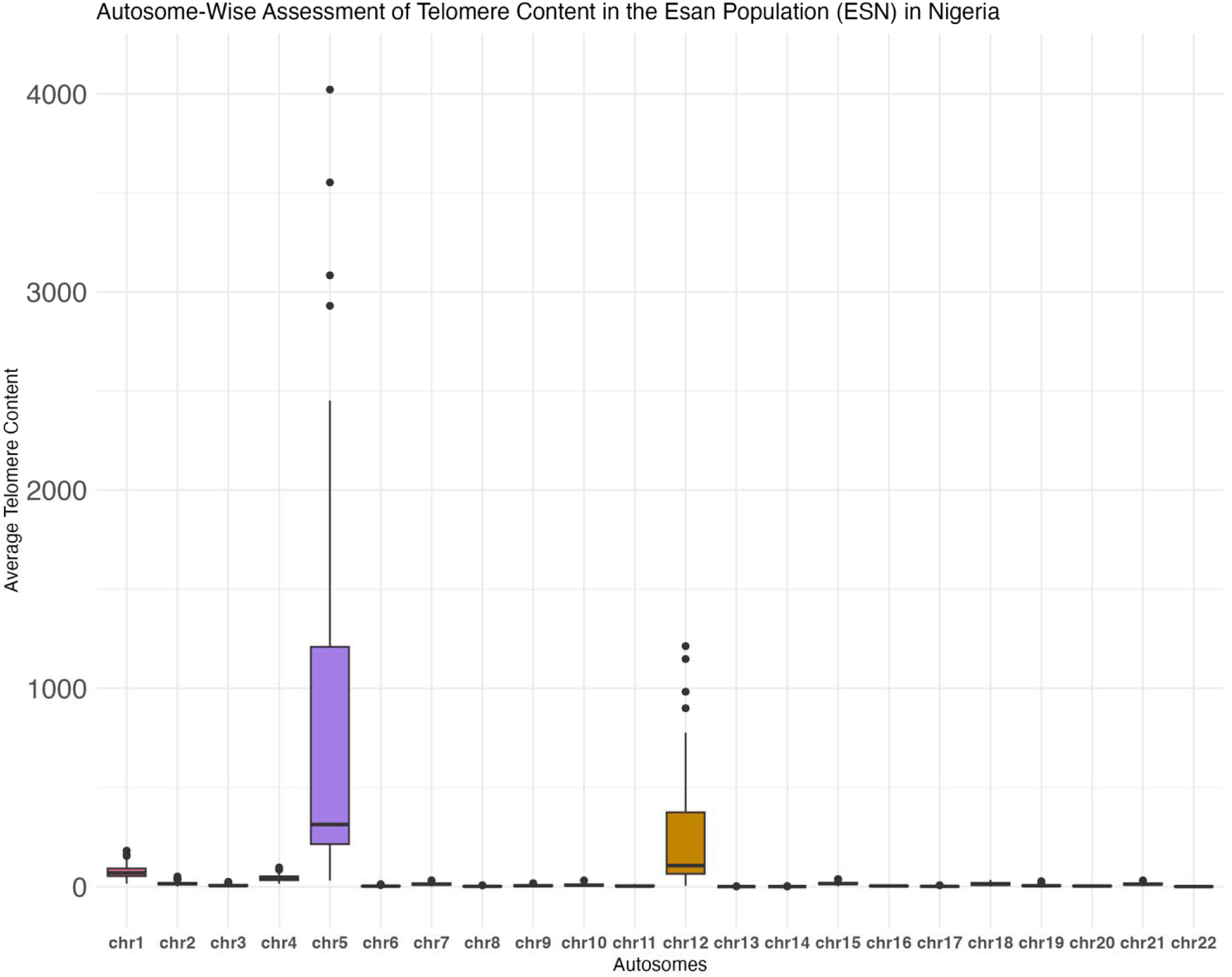
Autosome-wise telomere content distribution in the Esan Population in Nigeria (ESN). Box-and-whisker plot of autosome-wise assessment of average telomere content per autosome in ESN (n = 98).

**Fig 16.**
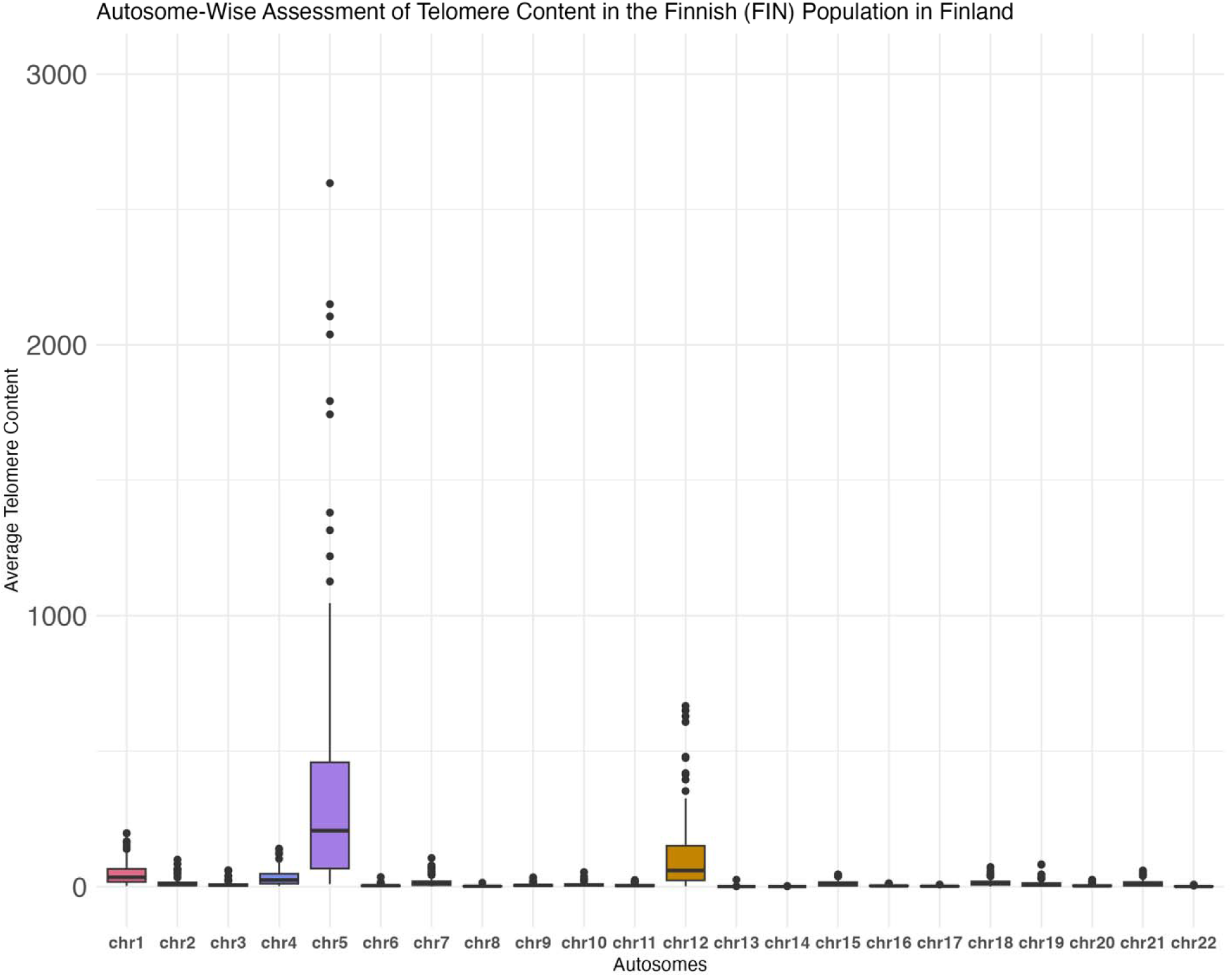
Autosome-wise telomere content distribution in the Finnish population in Finland (FIN). Box-and-whisker plot of autosome-wise assessment of average telomere content per autosome in FIN (n = 92).

**Fig 17.**
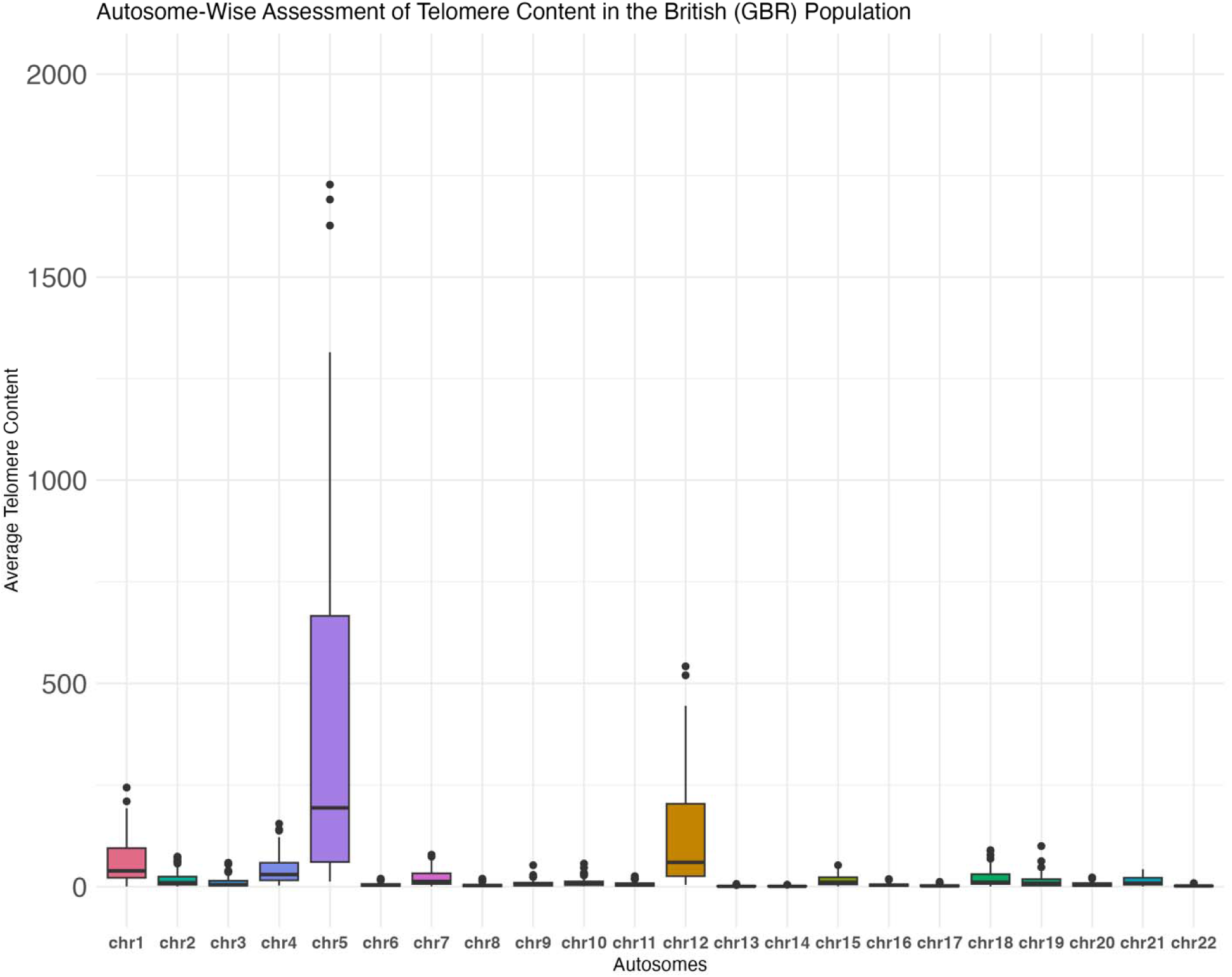
Autosome-wise telomere content distribution in the British population in the Great Britain (GBR). Box-and-whisker plot of autosome-wise assessment of average telomere content per autosome in GBR (n = 89).

**Fig 18.**
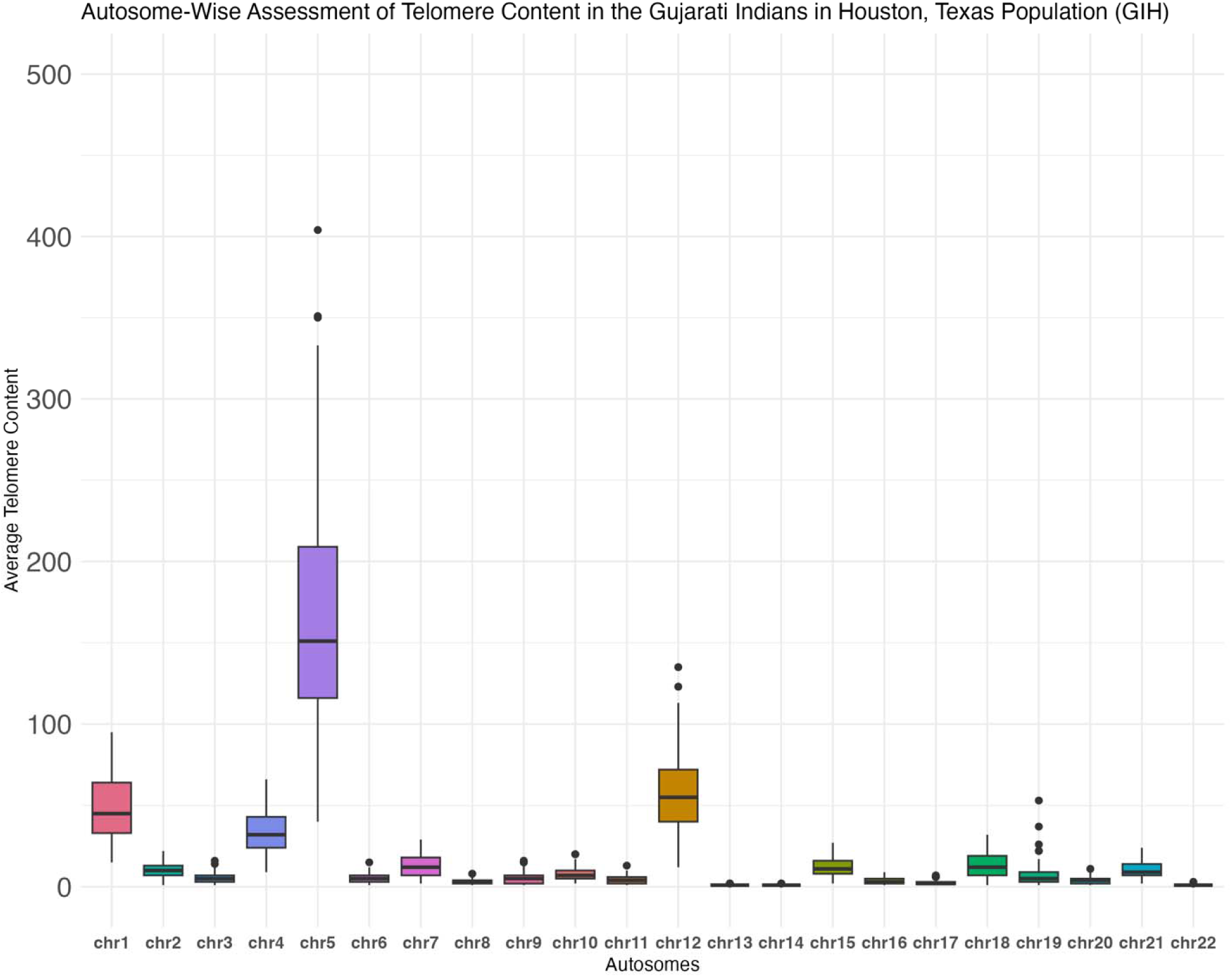
Autosome-wise telomere content distribution in the Gujarati Indian population in Houston, Texas (GIH). Box-and-whisker plot of autosome-wise assessment of average telomere content per autosome in GIH (n = 97).

**Fig 19.**
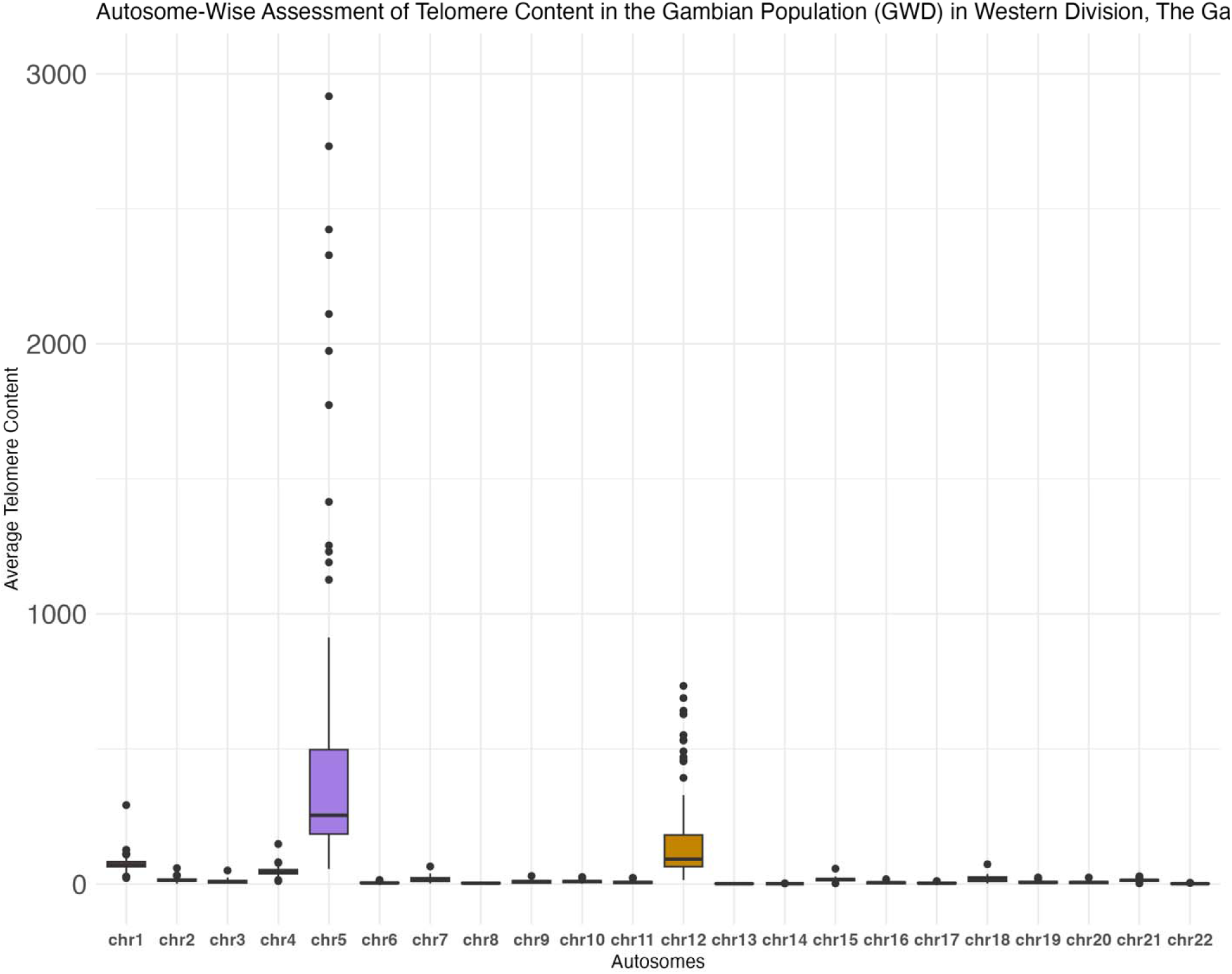
Autosome-wise telomere content distribution in the Gambian population in Western Division, the Gambia (GWD). Box-and-whisker plot of autosome-wise assessment of average telomere content per autosome in GWD (n = 110).

**Fig 20.**
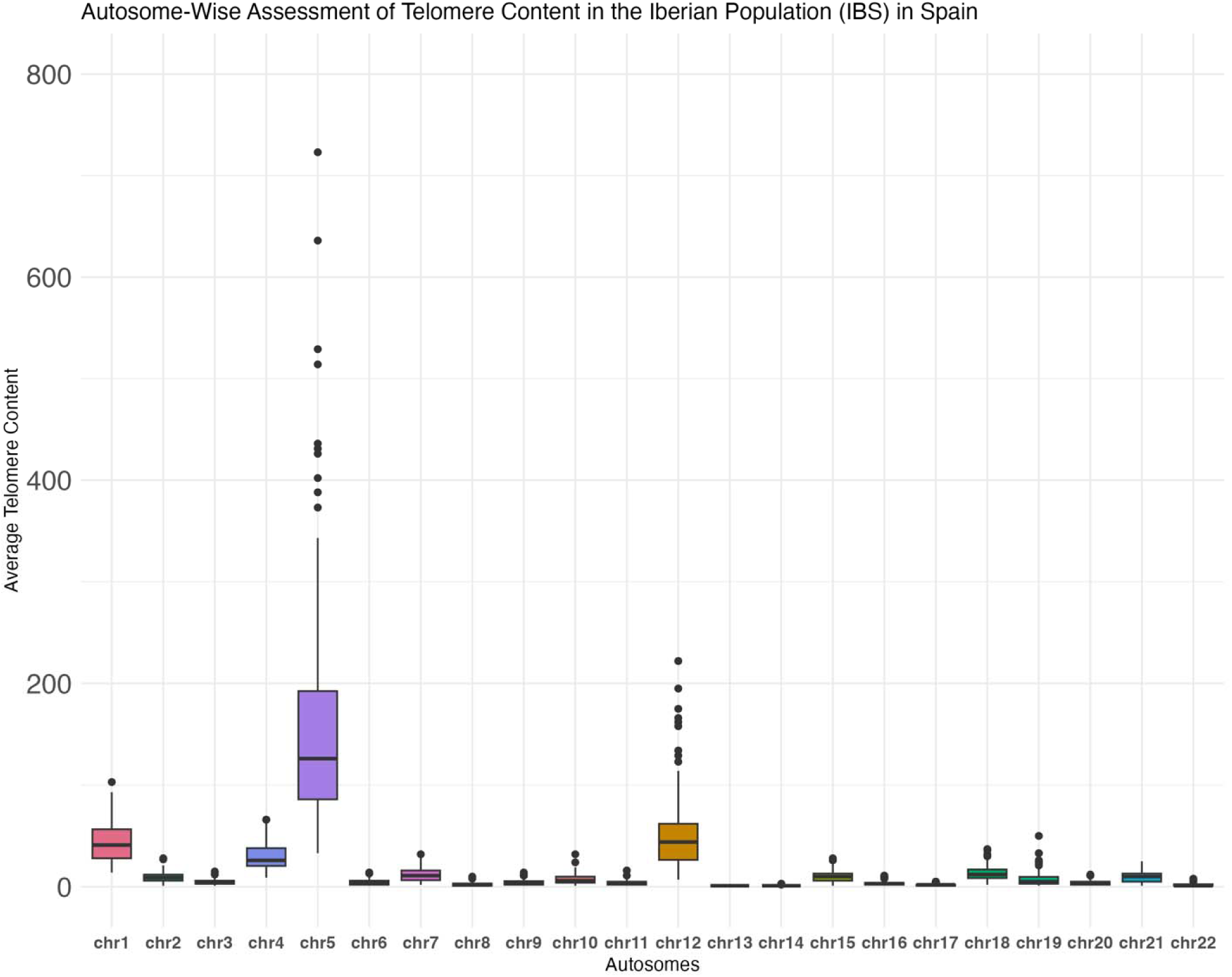
Autosome-wise telomere content distribution in the Iberian population in Spain (IBS). Box-and-whisker plot of autosome-wise assessment of average telomere content per autosome in IBS (n = 106).

**Fig 21.**
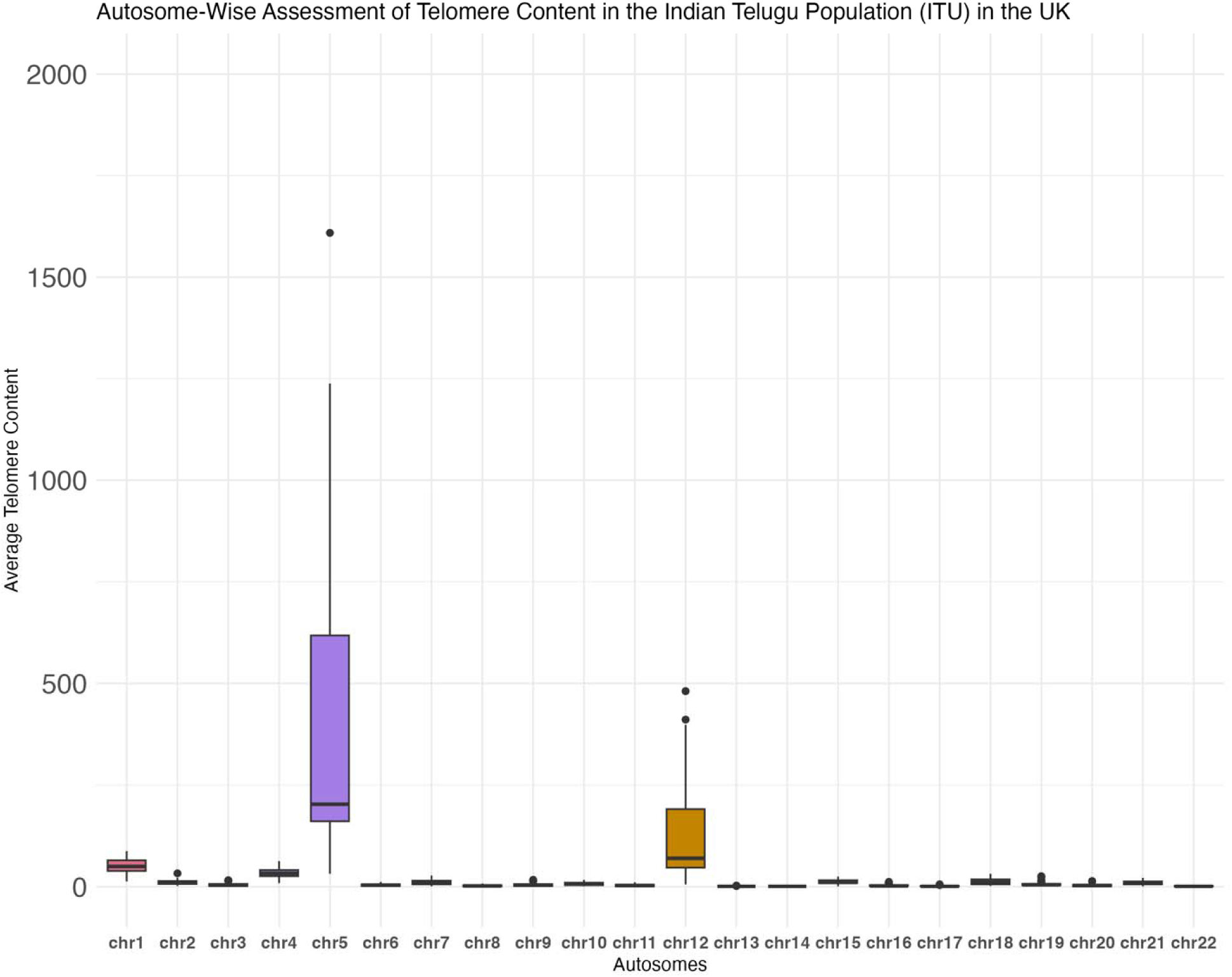
Autosome-wise telomere content distribution in the Indian Telugu population in the United Kingdom (ITU). Box-and-whisker plot of autosome-wise assessment of average telomere content per autosome in ITU (n = 89).

**Fig 22.**
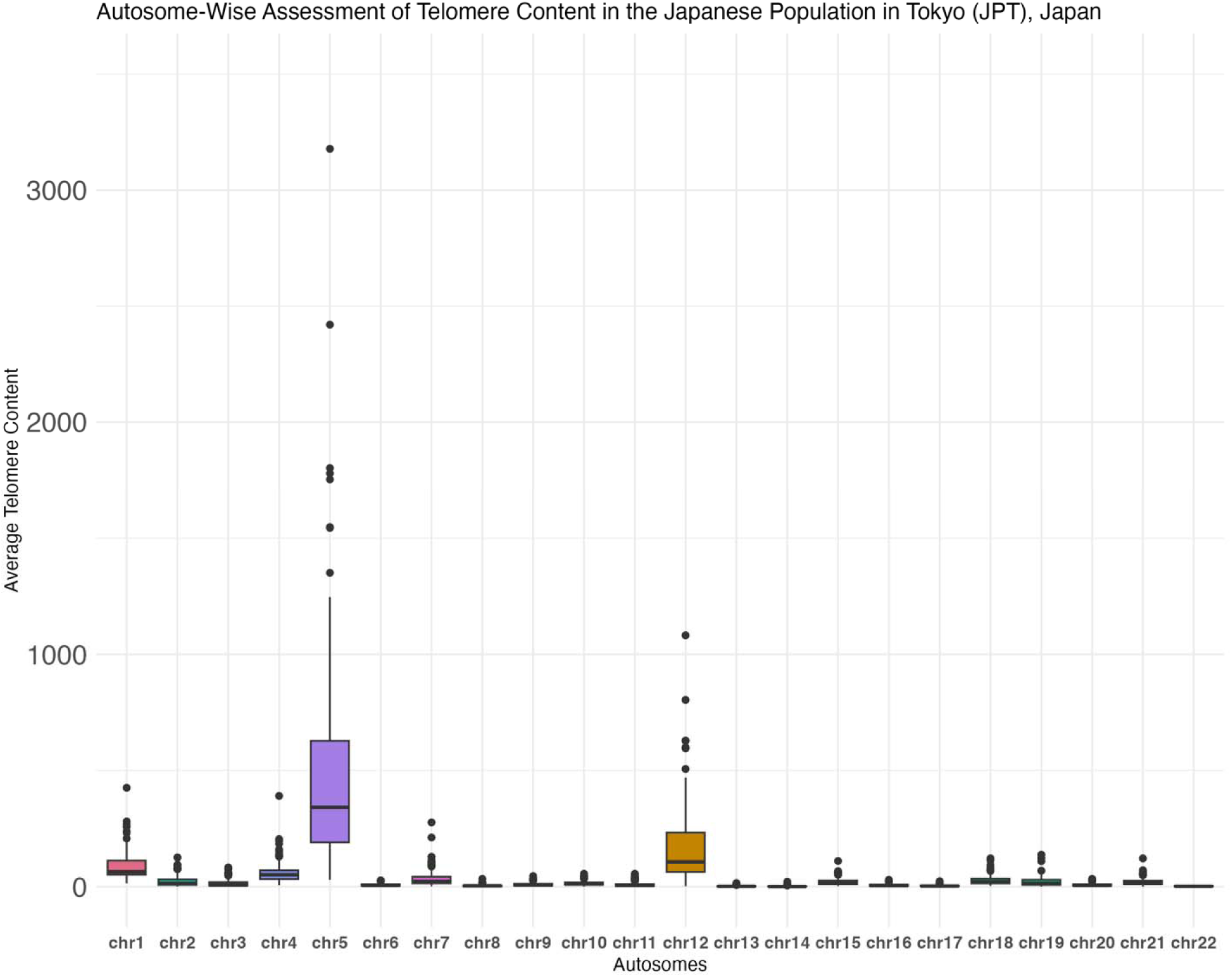
Autosome-wise telomere content distribution in the Japanese population in Tokyo, Japan (JPT). Box-and-whisker plot of autosome-wise assessment of average telomere content per autosome in JPT (n = 104).

**Fig 23.**
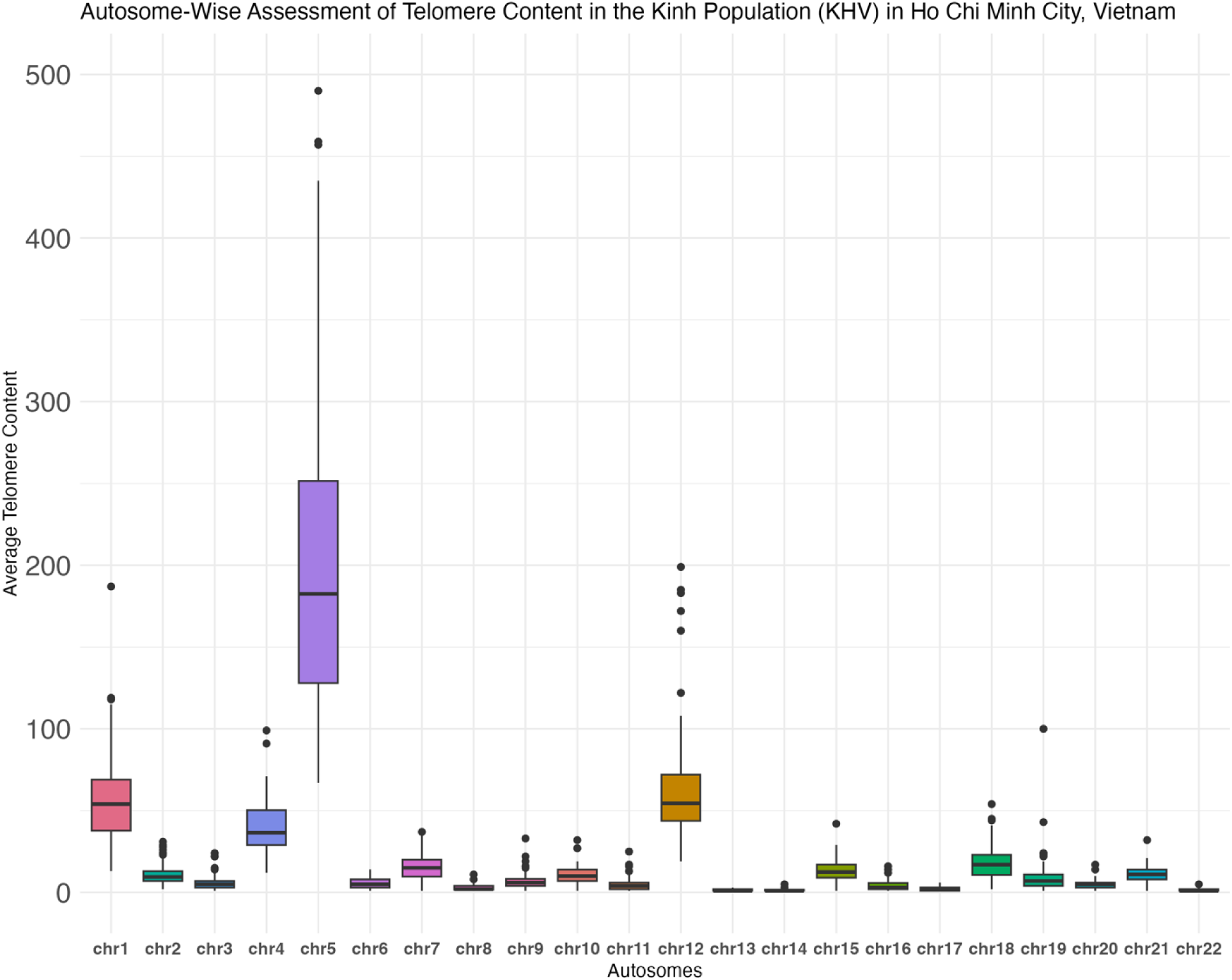
Autosome-wise telomere content distribution in the Kinh population in Ho Chi Minh City, Vietnam (KHV). Box-and-whisker plot of autosome-wise assessment of average telomere content per autosome in KHV (n = 88).

**Fig 24.**
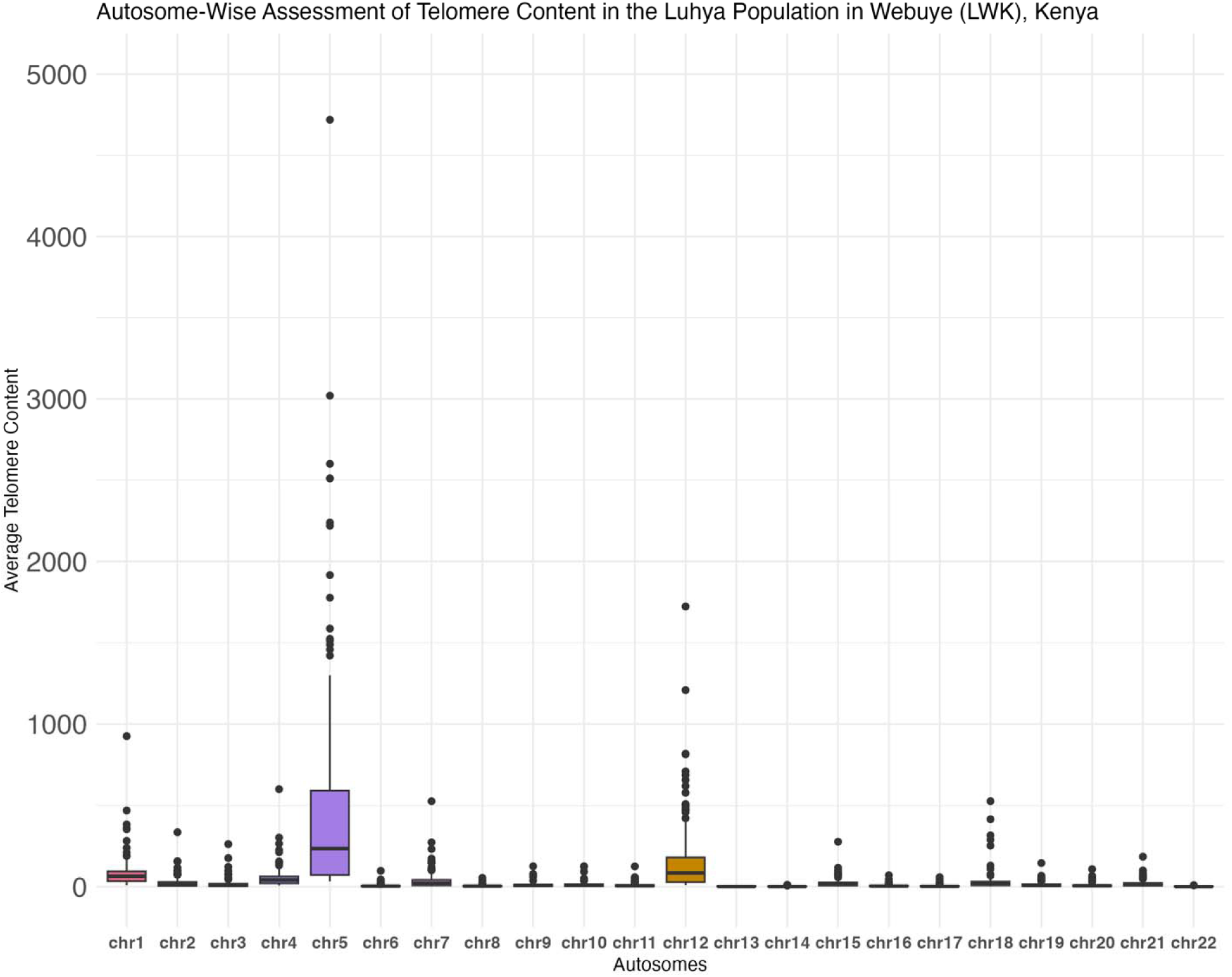
Autosome-wise telomere content distribution in the Luhya population in Webuye, Kenya (LWK). Box-and-whisker plot of autosome-wise assessment of average telomere content per autosome in LWK (n = 99).

**Fig 25.**
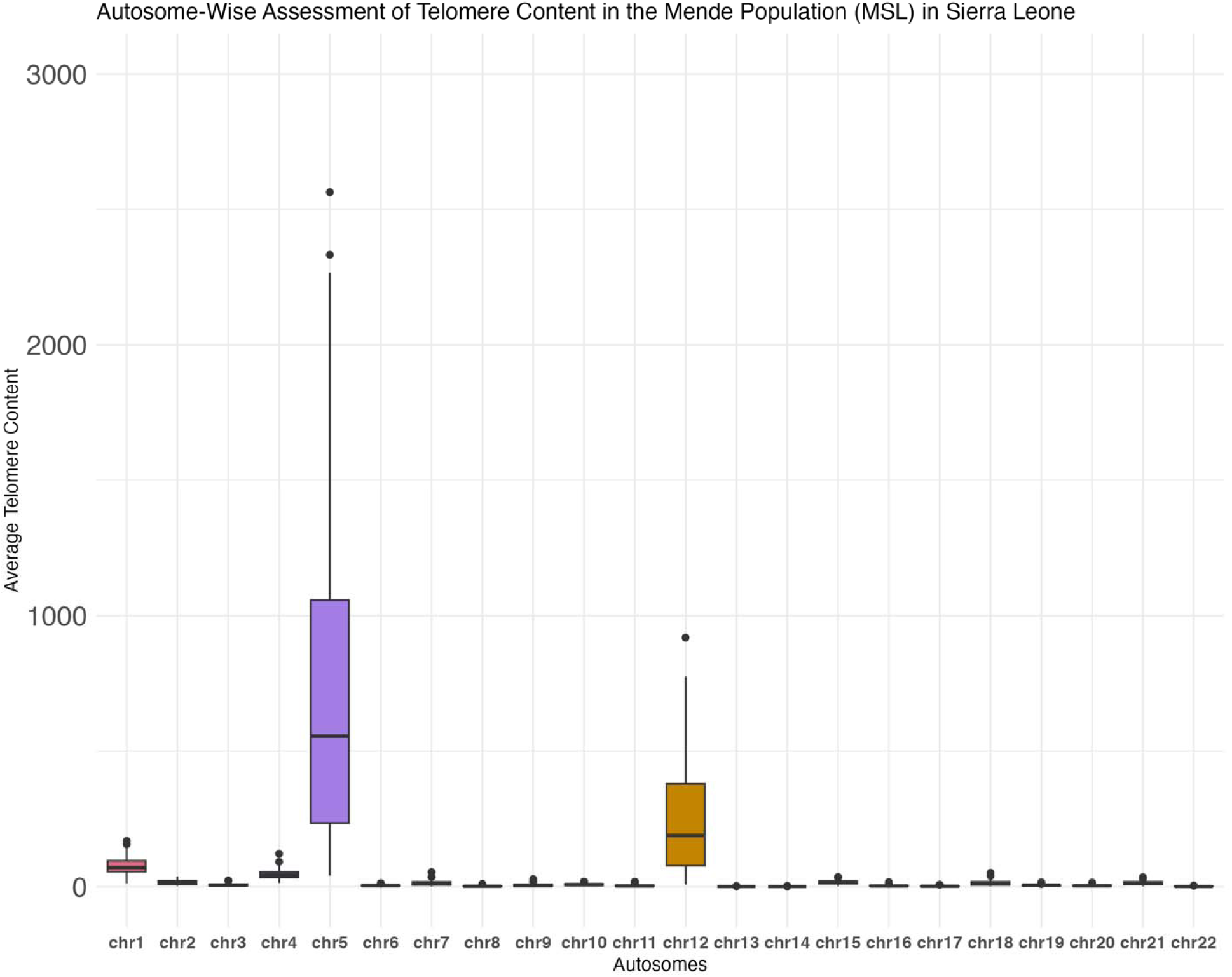
Autosome-wise telomere content distribution in the Mende population in Sierra Leone (MSL). Box-and-whisker plot of autosome-wise assessment of average telomere content per autosome in MSL (n = 84).

**Fig 26.**
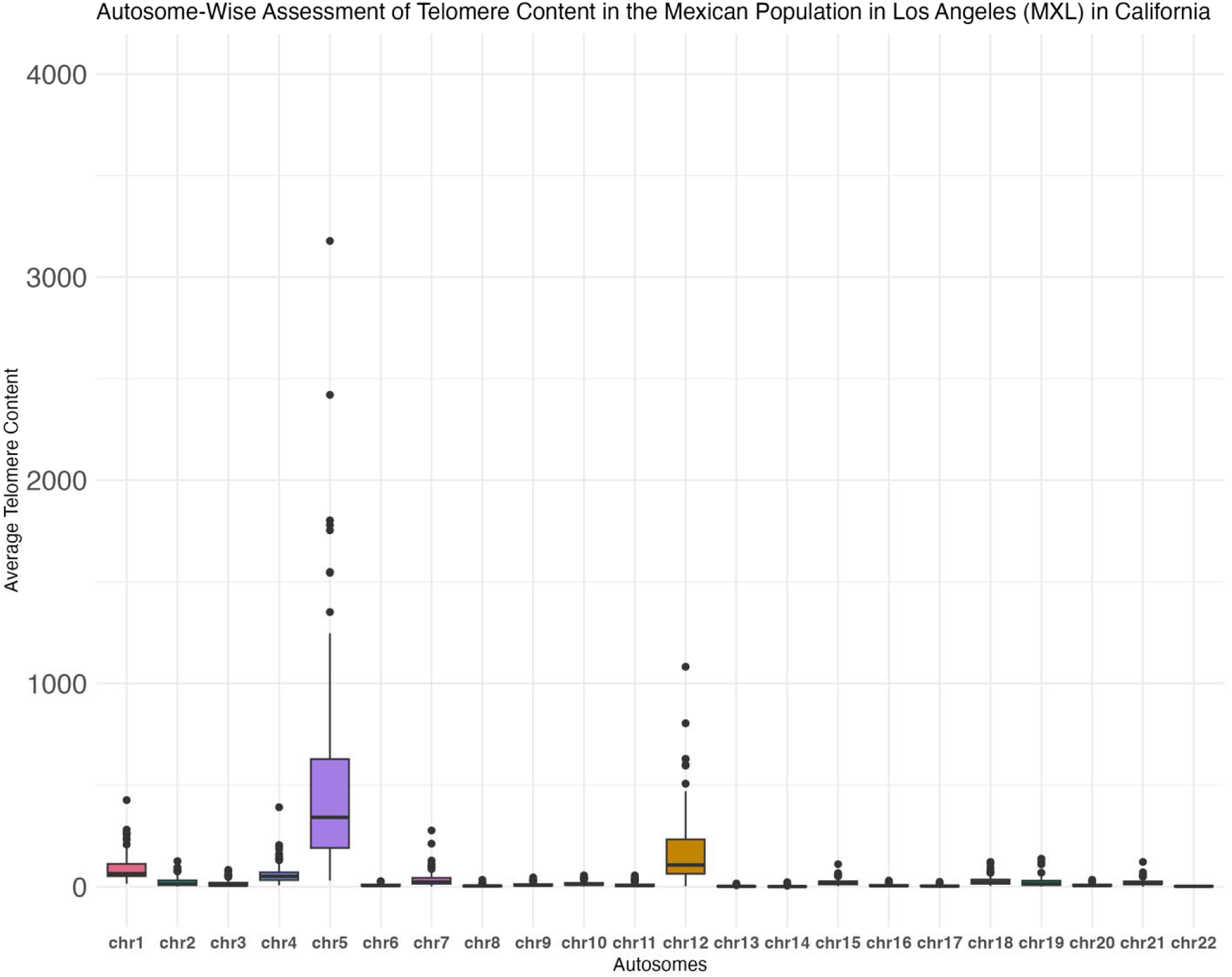
Autosome-wise telomere content distribution in the Mexican population in Los Angeles, California (MXL). Box-and-whisker plot of autosome-wise assessment of average telomere content per autosome in MXL (n = 66).

**Fig 27.**
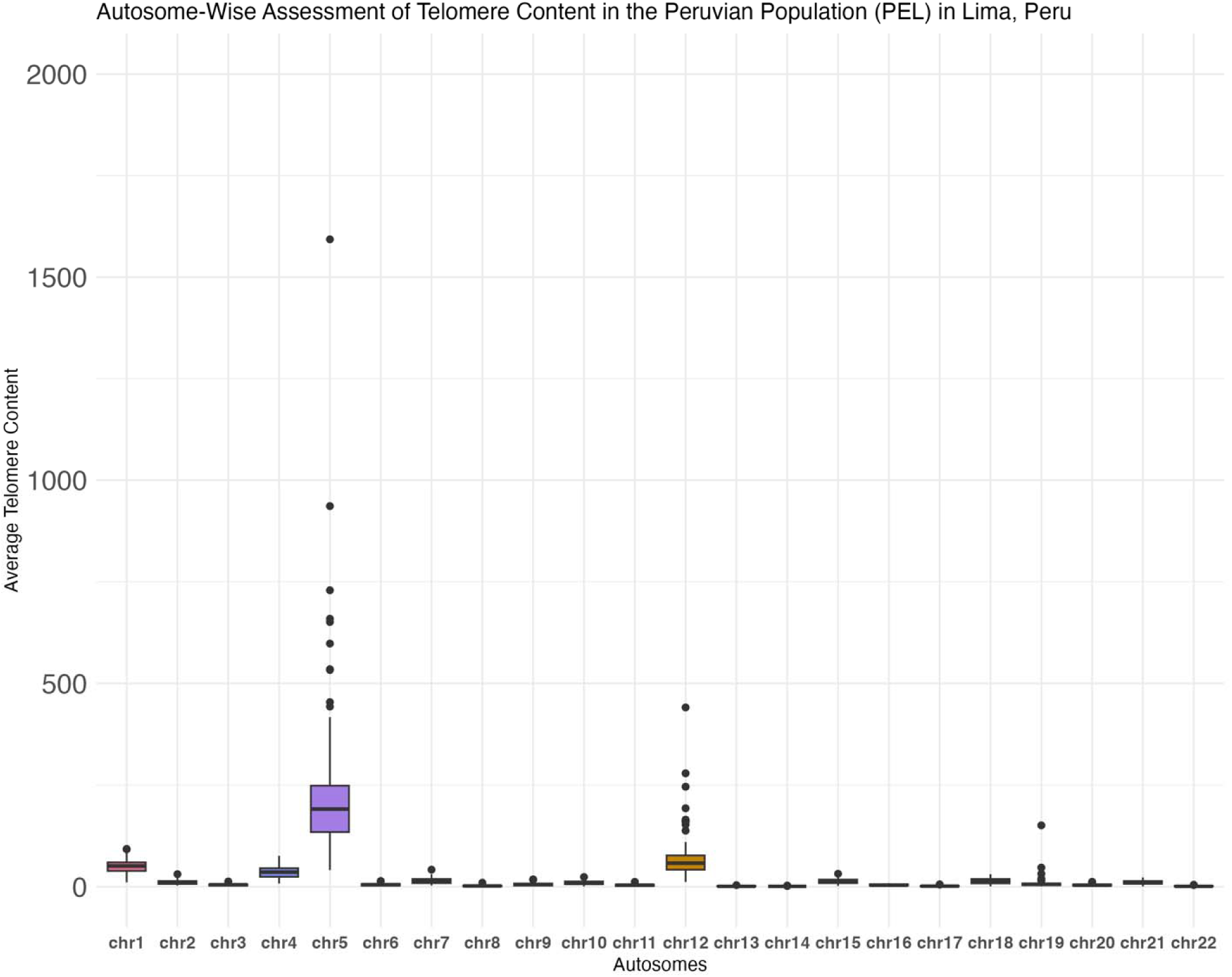
Autosome-wise telomere content distribution in the Peruvian population in Lima, Peru (PEL). Box-and-whisker plot of autosome-wise assessment of average telomere content per autosome in PEL (n = 83).

**Fig 28.**
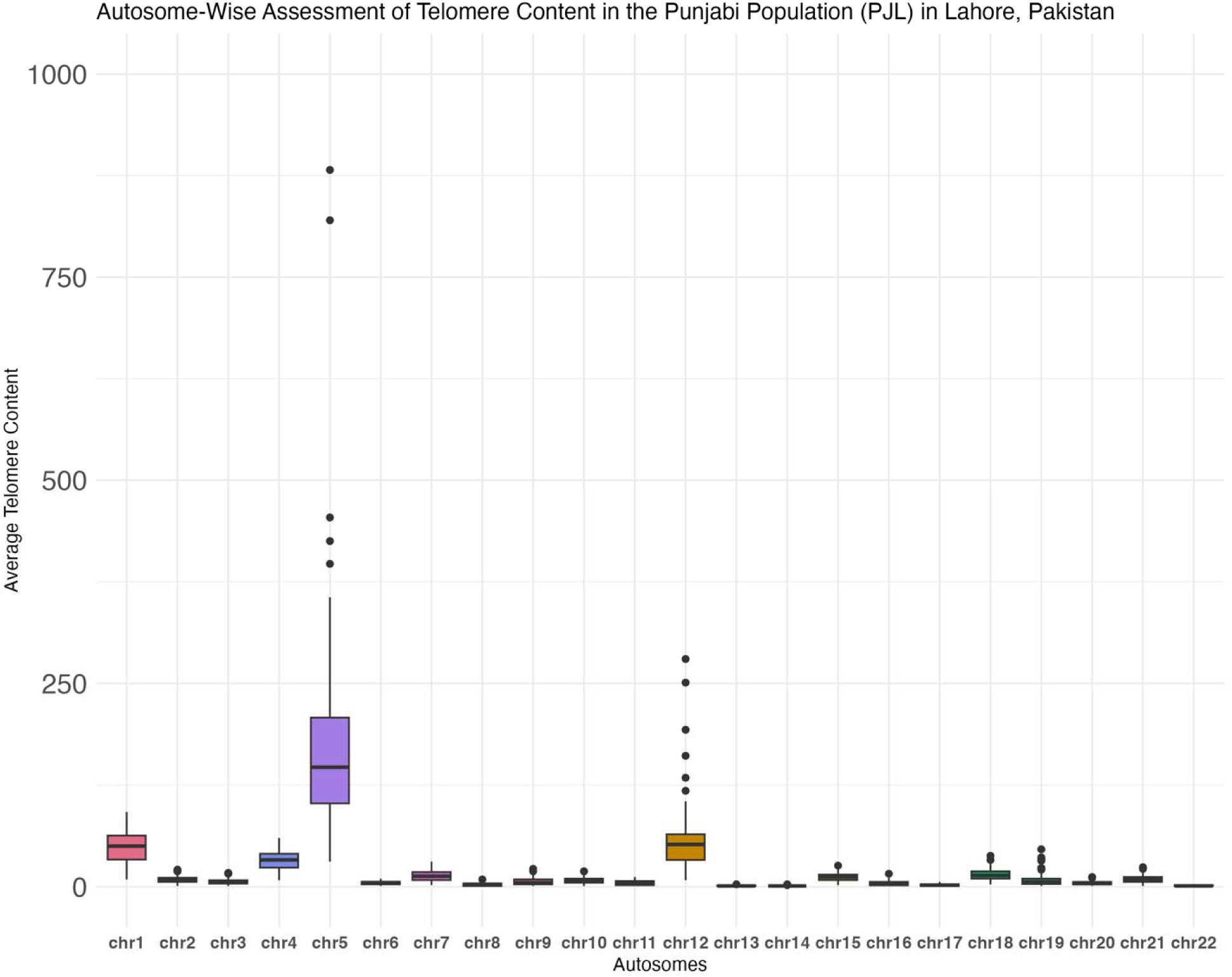
Autosome-wise telomere content distribution in the Punjabi population in Lahore, Pakistan (PJL). Box-and-whisker plot of autosome-wise assessment of average telomere content per autosome in PJL (n = 87).

**Fig 29.**
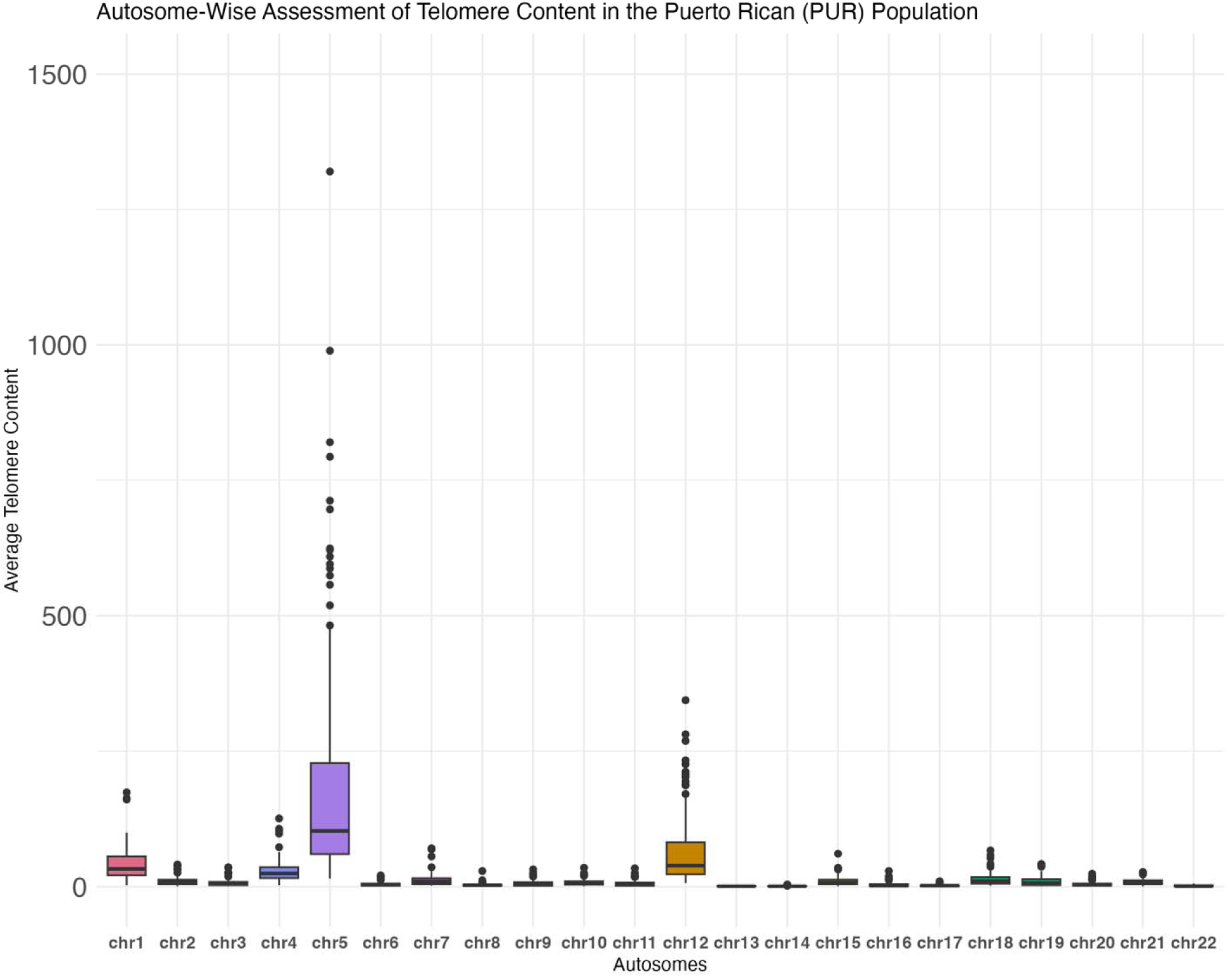
Autosome-wise telomere content distribution in the Puerto Rican population in Puerto Rico (PUR). Box-and-whisker plot of autosome-wise assessment of average telomere content per autosome in PUR (n = 102).

**Fig 30.**
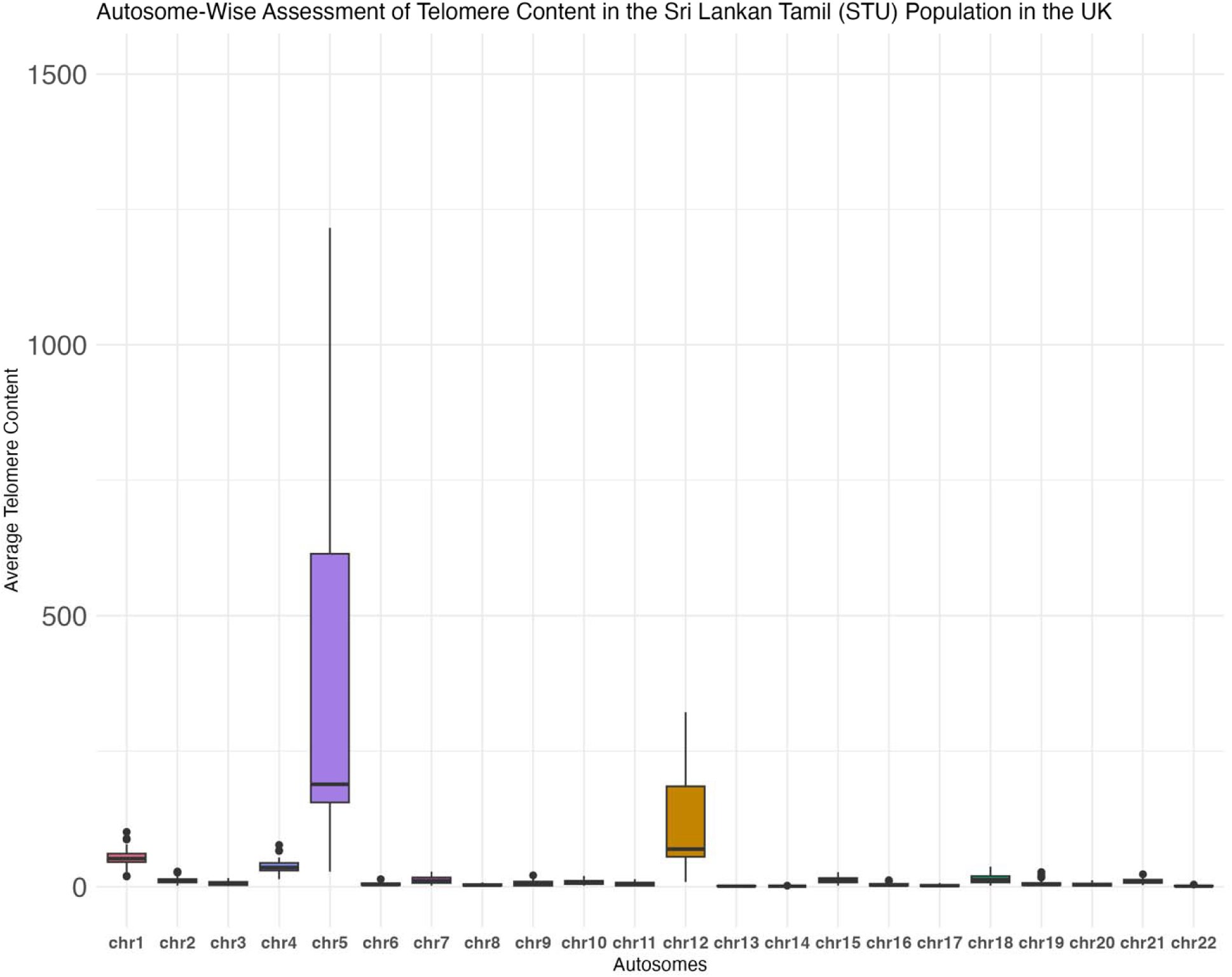
Autosome-wise telomere content distribution in the Sri Lankan Tamil population in the United Kingdom (STU). Box-and-whisker plot of autosome-wise assessment of average telomere content per autosome in STU (n = 74).

**Fig 31.**
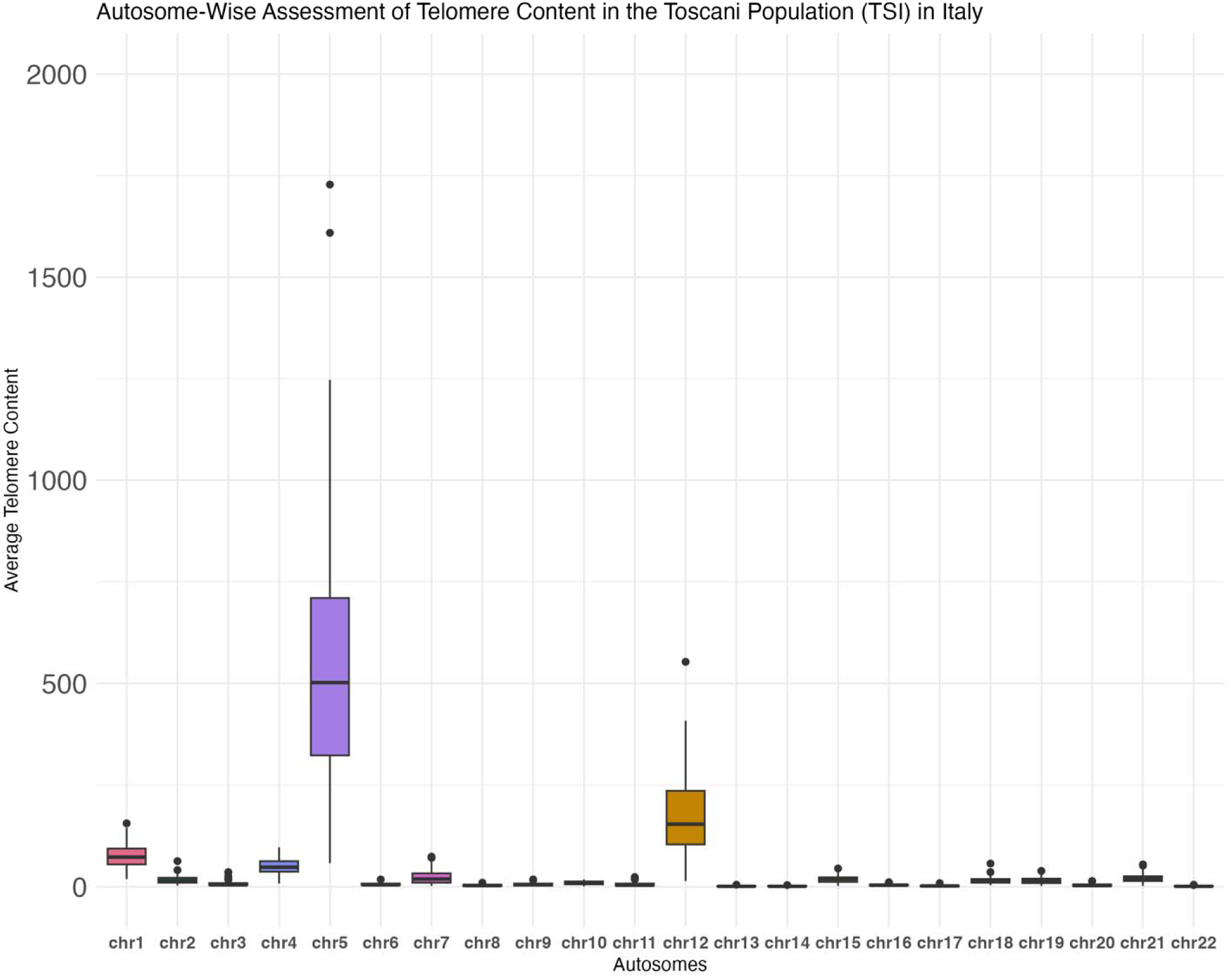
Autosome-wise telomere content distribution in the Toscani population in Italy (TSI). Box-and-whisker plot of autosome-wise assessment of average telomere content per autosome in TSI (n = 66).

**Fig 32.**
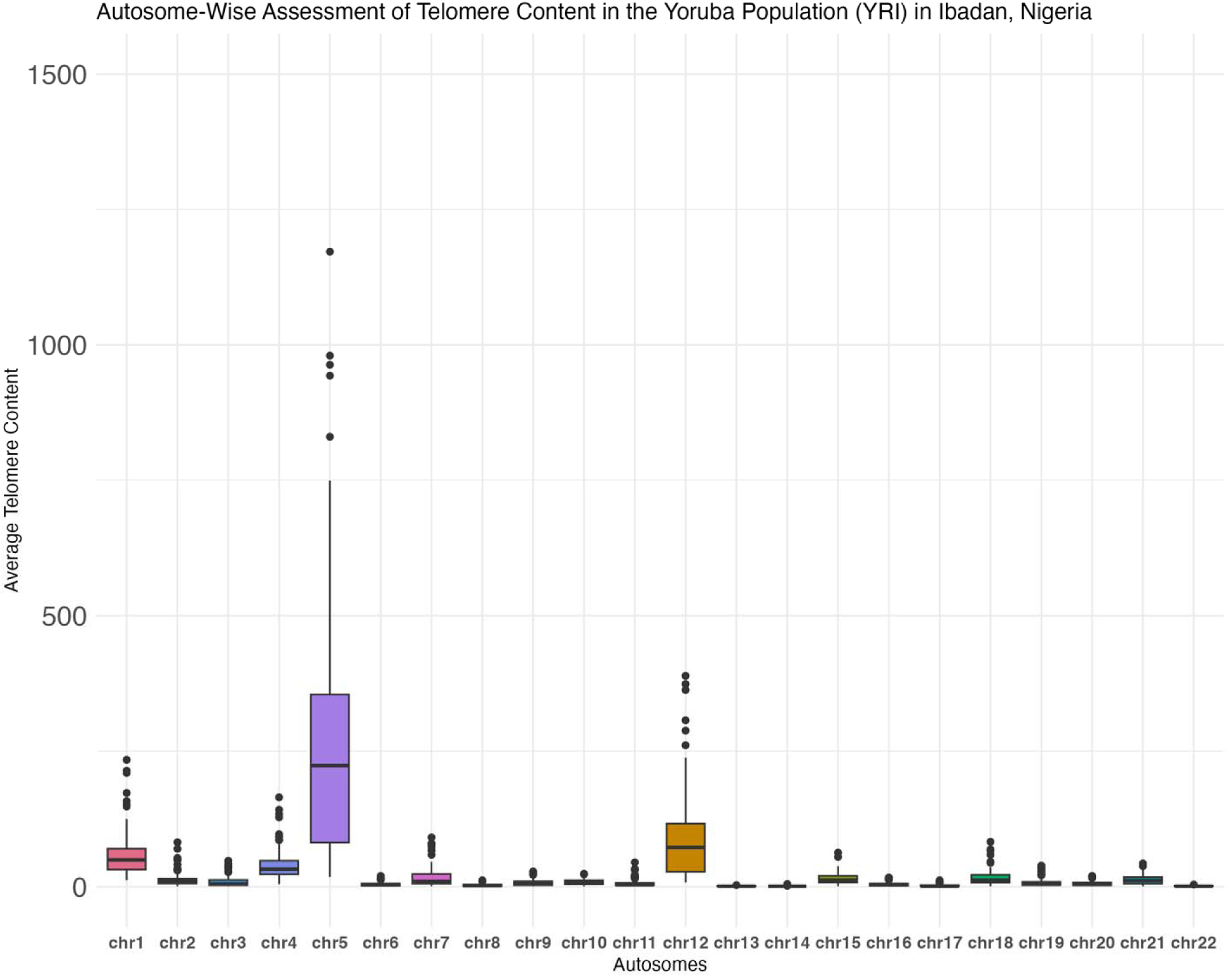
Autosome-wise telomere content distribution in the Yoruba population in Ibadan, Nigeria (YRI). Box-and-whisker plot of autosome-wise assessment of average telomere content per autosome in YRI (n = 108).

## DISCUSSION

Telomere content and length variation has been implicated in numerous complex human phenotypes, including the progression of senescence and cellular aging, cancers (The Telomeres Mendelian Randomization Collaboration et al., 2017), apoptosis, inflammatory response (Shammas, 2011), dementia and mortality (Grodstein et al., 2008; Honig et al., 2012). Previous studies have shown variation in all these phenotypes across human populations (Demanelis et al., 2020), however, no study as yet has collated and cataloged telomere variation and correlated it with phenotypic variation across global populations. As a first step towards this, here we catalog telomere content variation across 26 global populations using whole genomic sequence data collated from the 1000 Genomes Project. We develop a pipeline that incorporates a fast and memory efficient telomere content estimator, qmotif v.1.0 with the 1000 Genomes Project data, and implement time/memory reduction strategies (e.g. including only telomeric regions already known from the hg19 human reference genome, incorporating both small, and complex regular expressions) to build the first catalog of telomere content variation across modern humans.

Statistical analyses of this catalog showed (1) significant variation in telomere content among 26 populations, (2) significant differences in telomere content among superpopulations, (3) no significant differences in telomere content among self-reported sexes across the globe, or within populations, (4) consistently higher median and variability in telomere content across chromosomes 5 and 12 in all 26 populations.

While our study primarily serves to catalog variation in telomere content across modern humans, patterns of non-significant variation among self-reported sexes, within and between populations, and across superpopulations indicates that we are yet to understand the complexities of genomic and environmental variation that contributes to differences in longevity in sexes. Similar results were also observed in previous studies (Shammas, 2011) where summary estimates from only TRF Southern Blot method showed longer telomeres in females than males but neither Realtime PCR nor the Flow- FISH method results. A number of biological plausible arguments have been made for these sex-based differences in longevity. These have been proposed to be explained by behavioral differences in developed nations, child mortality rates (Zarulli & Salinari, 2024), rates of reproductive decline (Vinicius et al., 2014), pharmacological interventions (Austad & Bartke, 2016), response to inflammatory activity (Aviv, 2002), estrogen and other hormonal receptors (Aviv, 2002), and several other complex biological processes. The heterogametic sex hypothesis (Barrett & Richardson, 2011) proposes that any harmful recessive alleles on the X chromosome in males (the heterogametic sex in humans, with XY chromosomes) lack a compensatory counterpart, which differs from females who may have a second X chromosome to provide compensation (Austad, 2006). Consequently, males may experience shorter telomeres due to the presence of less effective telomere maintenance alleles on their unprotected X chromosome (Barrett & Richardson, 2011). In addition, having two X chromosomes has been shown to provide a health benefit for X-linked disorders (Christensen et al., 2000). We surmise that a more complete picture of the role of telomere content variation in sex-based differences in longevity will emerge with Biobank scale datasets that collate high-quality whole genomic, phenotypic, behavioral, and environmental data from globally diverse populations.

Telomere length is heritable (Armanios & Blackburn, 2012) and is strictly maintained during life (Graakjaer et al., 2004). (Baird, 2008) proposed a two-level telomere length control, where first trans-acting factors determine the genome-wide telomere length for each individual, and then cis-acting mechanisms determine the relative-chromosome specific length distribution at each chromosome end. Our analysis of autosome-wise assessment of telomeric reads across 26 global human populations also showed that long chromosomes like chromosome 5 and chromosome 12 contained higher telomere content, while chromosome 13-15, chromosome 17, chromosome 21, and chromosome 22 were the shortest, and also contained lower telomere content (Karimian et al., 2024) (Vasa-Nicotera et al., 2005). The shortest chromosomes are also some of the acrocentric human chromosomes, indicating that there are potential relationships between chromosomal structure and telomere content.

We selected 1000 Genomes Project data because we wanted our catalog of telomere content to reflect the diversity of modern human populations. However, the 1000 Genomes Project only includes self-reported sex and race as phenotypic data for all self-declared healthy individuals over the age of 18. The absence of age data is one of the caveats in this study. Furthermore, we acknowledge potential sampling biases where the group of participants represented in the 1000 Genomes Project do not accurately present the population, causing telomere content variation to be skewed or biased. We propose that future research be powered to investigate whether specific chromosomes consistently have highest or lowest telomere content throughout a wide age range.

Short telomere syndromes are genetically predisposed and worsen with each generation (Armanios & Blackburn, 2012). In order to make this catalog of telomere content variation clinically meaningful, we also propose that future studies should look at longitudinal measures and multi-generational measures of both telomere length and aging-related parameters to understand inheritance of telomere length. Furthermore, with each reference genome version, there is a progressive improvement in quality, which is made possible by technological breakthroughs and improved sequencing techniques, as well as an improvement in the reference genome’s representation of historically underrepresented populations. Since the reference genome we used (GRCh37/hg19) was originally produced utilizing donor samples from specific populations or restricted to certain geographical locations, we propose that efforts should be made to incorporate more sequences from diverse populations across various geographical locations in order to create an adequate representative reference genome that sufficiently captures variation in telomere content.

Most studies on telomere length and the changes in telomere dynamics throughout an individual’s life have traditionally focused on human diseases. Recently, there has been increasing interest in telomere biology among researchers outside of the fields of cell biology, human health, and epidemiology. Scientists in the fields of evolution, physiology, and ecology are now exploring variations in telomere dynamics across different species, the relationship between telomere loss and life-history traits, and how environmental conditions influence these factors (Nussey et al., 2014). This shift illustrates the expanding scope of telomere research beyond just human health.

**Table 1.** Five superpopulations (largely grouped by subcontinental origin) analyzed for telomere content variation with their population code on the 1000 Genomes Project database.

| No. | Description | Superpopulation Code |
| --- | --- | --- |
| 1. | East Asian | EAS |
| 2. | South Asian | SAS |
| 3. | African | AFR |
| 4. | European | EUR |
| 5. | American | AMR |

**Table 2.** Whole genomic data and population of origin code of each population available in phase 3 of the 1000 Genomes Project database analyzed for telomere content variation in this study.

| No. | Populations | Population code | Samples |
| --- | --- | --- | --- |
| 1. | African Caribbean in Barbados | ACB | 121 |
| 2. | African Ancestry in Southwest USA | ASW | 111 |
| 3. | Bengali in Bangladesh | BEB | 88 |
| 4. | British in England and Scotland | GBR | 104 |
| 5. | Utah residents (CEPH) with Northern and Western European ancestry | CEU | 183 |
| 6. | Chinese Dai in Xishuangbanna, China | CDX | 102 |
| 7. | Colombian in Medellin, Colombia | CLM | 132 |
| 8. | Esan in Nigeria | ESN | 100 |
| 9. | Finnish in Finland | FIN | 103 |
| 10. | Gambian in Western Division - Mandinka | GWD | 102 |
| 11. | Gujarati Indians in Houston, Texas | GIH | 113 |
| 12. | Han Chinese in Beijing, China | CHB | 108 |
| 13. | Han Chinese South | CHS | 165 |
| 14. | Iberian Populations in Spain | IBS | 157 |
| 15. | Indian Telugu in the UK | ITU | 103 |
| 16. | Japanese in Tokyo, Japan | JPT | 105 |
| 17. | Kinh in Ho Chi Minh City, Vietnam | KHV | 121 |
| 18. | Luhya in Webuye, Kenya | LWK | 116 |
| 19. | Mende in Sierra Leone | MSL | 90 |
| 20. | Mexican Ancestry in Los Angeles,<br>California | MXL | 104 |
| 21. | Peruvian in Lima, Peru | PEL | 122 |
| 22. | Puerto Rican in Puerto Rico | PUR | 139 |
| 23. | Punjabi in Lahore, Pakistan | PJL | 113 |
| 24. | Sri Lankan Tamil in the UK | STU | 105 |
| 25. | Toscani in Italy | TSI | 112 |
| 26. | Yoruba in Ibadan, Nigeria | YRI | 186 |
|  | Total Populations = 26 | Total samples = 3105 |  |

**Table 3.** Total self-reported male and female samples analyzed from Phase 3 of the 1000 Genomes Project database.

| Sr. No. | Population code | Female | Male | Total |
| --- | --- | --- | --- | --- |
| 1. | ACB | 45 | 43 | 88 |
| 2. | ASW | 39 | 26 | 65 |
| 3. | BEB | 43 | 41 | 84 |
| 4. | CDX | 37 | 46 | 83 |
| 5. | CEU | 47 | 45 | 92 |
| 6. | CHB | 56 | 45 | 101 |
| 7. | CHS | 54 | 50 | 104 |
| 8. | CLM | 51 | 43 | 94 |
| 9. | ESN | 45 | 53 | 98 |
| 10. | FIN | 56 | 36 | 92 |
| 11. | GBR | 44 | 45 | 89 |
| 12. | GIH | 44 | 53 | 97 |
| 13. | GWD | 57 | 53 | 110 |
| 14. | IBS | 53 | 53 | 106 |
| 15. | ITU | 37 | 52 | 89 |
| 16. | JPT | 48 | 56 | 104 |
| 17. | KHV | 48 | 40 | 88 |
| 18. | LWK | 54 | 45 | 99 |
| 19. | MSL | 43 | 41 | 84 |
| 20. | MXL | 34 | 32 | 66 |
| 21. | PEL | 43 | 40 | 83 |
| 22. | PJL | 42 | 45 | 87 |
| 23. | PUR | 50 | 52 | 102 |
| 24. | STU | 37 | 37 | 74 |
| 25. | TSI | 32 | 34 | 66 |
| 26. | YRI | 57 | 51 | 108 |
|  | Total Populations = 26 | 1196 | 1157 | 2353 |

## ACKNOWLEDGEMENTS

We would like to extend our heartfelt gratitude to the individuals who consented to have their samples analyzed for the 1000 Genomes Project. Your contributions are the foundation of this research. We would also like to thank Dr. Kyle Hasenstab for his valuable recommendations on earlier versions of this manuscript.

## AUTHOR CONTRIBUTIONS

AS conceived the study, obtained funding, directed research, and edited the manuscript. PS performed all the research, designed and implemented the software pipelines, collated results, and wrote the manuscript.

## DATA AVAILABILITY

The pipeline developed to catalog telomere content variation can be accessed at (https://github.com/paribytes/TeloTales). Whole genomic data used in this catalog was obtained from NCBI’s server for Phase 3 of the 1000 Genomes Project, which is publicly available at (https://www.ncbi.nlm.nih.gov/projects/faspftp/1000genomes/).

---

**Table S1.** Chromosomal coordinates in the hg19 human reference genome utilized in the INCLUDES section of the qmotif v1.0 configuration file to capture only telomeric regions determined. Here, p represents the short arm of the chromosome and q is the long arm of the chromosome.

| Chromosome number/Arm | Chromosomal coordinates (sequence: start-stop) |
| --- | --- |
| chr1p | chr1:10001-12464 |
| chr1q | chr1:249237907-249240620 |
| chr2p | chr2:10001-12592 |
| chr2q | chr2:243187373-243189372 |
| chr2xA | chr2:243150480-243154648 |
| chr3p | chr3:60001-62000 |
| chr3q | chr3:197960430-197962429 |
| chr3xB | chr3:197897576-197903397 |
| chr4p | chr4:10001-12193 |
| chr4q | chr4:191041613-191044275 |
| chr5p | chr5:10001-13806 |
| chr5q | chr5:180903260-180905259 |
| chr6p | chr6:60001-62000 |
| chr6q | chr6:171053067-171055066 |
| chr7p | chr7:10001-12238 |
| chr7q | chr7:159126558-159128662 |
| chr8p | chr8:10001-12000 |
| chr8q | chr8:146302022-146304021 |
| chr9p | chr9:10001-12359 |
| chr9q | chr9:141151431-141153430 |
| chr10p | chr10:60001-62000 |
| chr10q | chr10:135522469-135524746 |
| chr11p | chr11:60001-62000 |
| chr11q | chr11:134944458-134946515 |
| chr12p | chr12:60001-62000 |
| chr12q | chr12:133839458-133841894 |
| chr12xC | chr12:93158-97735 |
| chr13p | chr13:19020001-19022000 |
| chr13q | chr13:115107878-115109877 |
| chr14p | chr14:19020001-19022000 |
| chr14q | chr14:107287540-107289539 |
| chr15p | chr15:20000001-20002000 |
| chr15q | chr15:102518969-102521391 |
| chr16p | chr16:60001-62033 |
| chr16q | chr16:90292753-90294752 |
| chr17p | chr17:1-2000 |
| chr17q | chr17:81193211-81195210 |
| chr18p | chr18:10001-12621 |
| chr18q | chr18:78014226-78017247 |
| chr19p | chr19:60001-62000 |
| chr19q | chr19:59116822-59118982 |
| chr20p | chr20:60001-62000 |
| chr20q | chr20:62963520-62965519 |
| chr21p | chr21:9411194-9413193 |
| chr21q | chr21:48117788-48119894 |
| chr22p | chr22:16050001-16052000 |
| chr22q | chr22:51242566-51244565 |
| chrXp | chrX:60001-62033 |
| chrXq | chrX:155257733-155260559 |
| chrYp | chrY:10001-12033 |
| chrYq | chrY:59360739-59363565 |

**Table S2.** Pairwise comparisons of telomere content variation between populations using Games Howell post-hoc test. Significant differences are highlighted in the table below.

| Population 1 | Population 2 | Adjusted p-value |
| --- | --- | --- |
| ACB | ASW | 1.04e-05 |
| ACB | BEB | 5.01e-07 |
| ACB | CDX | 9.98e-01 |
| ACB | CEU | 9.95e-01 |
| ACB | CHB | 9.88e-01 |
| ACB | CHS | 1.00e+00 |
| ACB | CLM | 1.00e+00 |
| ACB | ESN | 7.69e-05 |
| ACB | FIN | 1.00e+00 |
| ACB | GBR | 1.00e+00 |
| ACB | GIH | 1.00e+00 |
| ACB | GWD | 4.32e-01 |
| ACB | IBS | 8.36e-01 |
| ACB | ITU | 1.00e+00 |
| ACB | JPT | 1.00e-02 |
| ACB | KHV | 9.63e-01 |
| ACB | LWK | 8.02e-01 |
| ACB | MSL | 6.94e-07 |
| ACB | MXL | 4.99e-09 |
| ACB | PEL | 1.00e+00 |
| ACB | PJL | 3.00e-03 |
| ACB | PUR | 1.00e-02 |
| ACB | STU | 1.00e+00 |
| ACB | TSI | 6.08e-06 |
| ACB | YRI | 1.00e+00 |
| ASW | BEB | 7.17e-14 |
| ASW | CDX | 2.40e-09 |
| ASW | CEU | 8.00e-02 |
| ASW | CHB | 1.37e-09 |
| ASW | CHS | 8.73e-04 |
| ASW | CLM | 4.05e-04 |
| ASW | ESN | 1.45e-13 |
| ASW | FIN | 1.30e-02 |
| ASW | GBR | 1.87e-01 |
| ASW | GIH | 1.37e-06 |
| ASW | GWD | 5.41e-12 |
| ASW | IBS | 1.76e-01 |
| ASW | ITU | 1.44e-06 |
| ASW | JPT | 2.98e-13 |
| ASW | KHV | 8.89e-04 |
| ASW | LWK | 9.38e-07 |
| ASW | MSL | 5.85e-14 |
| ASW | MXL | 8.14e-01 |
| ASW | PEL | 6.74e-04 |
| ASW | PJL | 9.96e-01 |
| ASW | PUR | 1.00e+00 |
| ASW | STU | 6.57e-05 |
| ASW | TSI | 1.30e-13 |
| ASW | YRI | 4.16e-04 |
| BEB | CDX | 1.71e-04 |
| BEB | CEU | 1.52e-09 |
| BEB | CHB | 6.50e-04 |
| BEB | CHS | 1.68e-07 |
| BEB | CLM | 1.74e-05 |
| BEB | ESN | 1.00e+00 |
| BEB | FIN | 2.00e-03 |
| BEB | GBR | 8.18e-06 |
| BEB | GIH | 1.71e-09 |
| BEB | GWD | 2.80e-02 |
| BEB | IBS | 3.65e-11 |
| BEB | ITU | 2.00e-03 |
| BEB | JPT | 4.68e-01 |
| BEB | KHV | 3.88e-11 |
| BEB | LWK | 3.19e-01 |
| BEB | MSL | 1.00e+00 |
| BEB | MXL | 0.00e+00 |
| BEB | PEL | 2.37e-08 |
| BEB | PJL | 2.32e-13 |
| BEB | PUR | 3.06e-13 |
| BEB | STU | 7.34e-04 |
| BEB | TSI | 1.00e+00 |
| BEB | YRI | 5.37e-08 |
| CDX | CEU | 2.65e-01 |
| CDX | CHB | 1.00e+00 |
| CDX | CHS | 9.58e-01 |
| CDX | CLM | 1.00e+00 |
| CDX | ESN | 1.40e-02 |
| CDX | FIN | 1.00e+00 |
| CDX | GBR | 9.79e-01 |
| CDX | GIH | 5.29e-01 |
| CDX | GWD | 1.00e+00 |
| CDX | IBS | 3.50e-02 |
| CDX | ITU | 1.00e+00 |
| CDX | JPT | 5.21e-01 |
| CDX | KHV | 4.60e-02 |
| CDX | LWK | 1.00e+00 |
| CDX | MSL | 3.55e-04 |
| CDX | MXL | 1.99e-12 |
| CDX | PEL | 8.12e-01 |
| CDX | PJL | 1.56e-06 |
| CDX | PUR | 1.81e-05 |
| CDX | STU | 1.00e+00 |
| CDX | TSI | 3.00e-03 |
| CDX | YRI | 9.08e-01 |
| CEU | CHB | 1.74e-01 |
| CEU | CHS | 1.00e+00 |
| CEU | CLM | 9.97e-01 |
| CEU | ESN | 2.64e-07 |
| CEU | FIN | 9.99e-01 |
| CEU | GBR | 1.00e+00 |
| CEU | GIH | 1.00e+00 |
| CEU | GWD | 8.00e-03 |
| CEU | IBS | 1.00e+00 |
| CEU | ITU | 5.30e-01 |
| CEU | JPT | 3.48e-05 |
| CEU | KHV | 1.00e+00 |
| CEU | LWK | 1.06e-01 |
| CEU | MSL | 1.56e-09 |
| CEU | MXL | 1.28e-04 |
| CEU | PEL | 1.00e+00 |
| CEU | PJL | 8.40e-01 |
| CEU | PUR | 8.08e-01 |
| CEU | STU | 8.73e-01 |
| CEU | TSI | 1.25e-08 |
| CEU | YRI | 1.00e+00 |
| CHB | CHS | 8.89e-01 |
| CHB | CLM | 1.00e+00 |
| CHB | ESN | 4.10e-02 |
| CHB | FIN | 1.00e+00 |
| CHB | GBR | 9.43e-01 |
| CHB | GIH | 3.71e-01 |
| CHB | GWD | 1.00e+00 |
| CHB | IBS | 2.00e-02 |
| CHB | ITU | 1.00e+00 |
| CHB | JPT | 7.76e-01 |
| CHB | KHV | 2.60e-02 |
| CHB | LWK | 1.00e+00 |
| CHB | MSL | 1.00e-03 |
| CHB | MXL | 1.08e-12 |
| CHB | PEL | 6.70e-01 |
| CHB | PJL | 8.83e-07 |
| CHB | PUR | 9.62e-06 |
| CHB | STU | 1.00e+00 |
| CHB | TSI | 1.20e-02 |
| CHB | YRI | 8.01e-01 |
| CHS | CLM | 1.00e+00 |
| CHS | ESN | 2.59e-05 |
| CHS | FIN | 1.00e+00 |
| CHS | GBR | 1.00e+00 |
| CHS | GIH | 1.00e+00 |
| CHS | GWD | 2.16e-01 |
| CHS | IBS | 9.97e-01 |
| CHS | ITU | 9.90e-01 |
| CHS | JPT | 3.00e-03 |
| CHS | KHV | 1.00e+00 |
| CHS | LWK | 5.82e-01 |
| CHS | MSL | 2.40e-07 |
| CHS | MXL | 5.25e-07 |
| CHS | PEL | 1.00e+00 |
| CHS | PJL | 8.40e-02 |
| CHS | PUR | 1.13e-01 |
| CHS | STU | 1.00e+00 |
| CHS | TSI | 2.14e-06 |
| CHS | YRI | 1.00e+00 |
| CLM | ESN | 2.00e-03 |
| CLM | FIN | 1.00e+00 |
| CLM | GBR | 1.00e+00 |
| CLM | GIH | 1.00e+00 |
| CLM | GWD | 8.35e-01 |
| CLM | IBS | 9.15e-01 |
| CLM | ITU | 1.00e+00 |
| CLM | JPT | 1.05e-01 |
| CLM | KHV | 9.88e-01 |
| CLM | LWK | 9.47e-01 |
| CLM | MSL | 3.77e-05 |
| CLM | MXL | 3.34e-07 |
| CLM | PEL | 1.00e+00 |
| CLM | PJL | 3.20e-02 |
| CLM | PUR | 4.10e-02 |
| CLM | STU | 1.00e+00 |
| CLM | TSI | 3.36e-04 |
| CLM | YRI | 1.00e+00 |
| ESN | FIN | 4.60e-02 |
| ESN | GBR | 5.45e-04 |
| ESN | GIH | 2.65e-07 |
| ESN | GWD | 4.78e-01 |
| ESN | IBS | 6.35e-09 |
| ESN | ITU | 6.40e-02 |
| ESN | JPT | 9.95e-01 |
| ESN | KHV | 5.19e-09 |
| ESN | LWK | 9.36e-01 |
| ESN | MSL | 1.00e+00 |
| ESN | MXL | 1.40e-13 |
| ESN | PEL | 4.03e-06 |
| ESN | PJL | 3.76e-13 |
| ESN | PUR | 1.64e-12 |
| ESN | STU | 3.20e-02 |
| ESN | TSI | 1.00e+00 |
| ESN | YRI | 9.12e-06 |
| FIN | GBR | 1.00e+00 |
| FIN | GIH | 1.00e+00 |
| FIN | GWD | 9.95e-01 |
| FIN | IBS | 9.70e-01 |
| FIN | ITU | 1.00e+00 |
| FIN | JPT | 6.09e-01 |
| FIN | KHV | 9.97e-01 |
| FIN | LWK | 9.98e-01 |
| FIN | MSL | 4.00e-03 |
| FIN | MXL | 5.85e-05 |
| FIN | PEL | 1.00e+00 |
| FIN | PJL | 2.01e-01 |
| FIN | PUR | 1.80e-01 |
| FIN | STU | 1.00e+00 |
| FIN | TSI | 2.50e-02 |
| FIN | YRI | 1.00e+00 |
| GBR | GIH | 1.00e+00 |
| GBR | GWD | 4.43e-01 |
| GBR | IBS | 1.00e+00 |
| GBR | ITU | 9.92e-01 |
| GBR | JPT | 3.40e-02 |
| GBR | KHV | 1.00e+00 |
| GBR | LWK | 6.64e-01 |
| GBR | MSL | 1.97e-05 |
| GBR | MXL | 2.00e-03 |
| GBR | PEL | 1.00e+00 |
| GBR | PJL | 8.44e-01 |
| GBR | PUR | 7.87e-01 |
| GBR | STU | 1.00e+00 |
| GBR | TSI | 1.63e-04 |
| GBR | YRI | 1.00e+00 |
| GIH | GWD | 1.30e-02 |
| GIH | IBS | 9.72e-01 |
| GIH | ITU | 8.68e-01 |
| GIH | JPT | 1.95e-05 |
| GIH | KHV | 9.99e-01 |
| GIH | LWK | 2.74e-01 |
| GIH | MSL | 8.95e-10 |
| GIH | MXL | 1.38e-09 |
| GIH | PEL | 1.00e+00 |
| GIH | PJL | 1.00e-03 |
| GIH | PUR | 1.30e-02 |
| GIH | STU | 9.95e-01 |
| GIH | TSI | 5.74e-09 |
| GIH | YRI | 1.00e+00 |
| GWD | IBS | 3.70e-04 |
| GWD | ITU | 1.00e+00 |
| GWD | JPT | 1.00e+00 |
| GWD | KHV | 3.10e-04 |
| GWD | LWK | 1.00e+00 |
| GWD | MSL | 6.50e-02 |
| GWD | MXL | 2.07e-13 |
| GWD | PEL | 7.70e-02 |
| GWD | PJL | 3.69e-09 |
| GWD | PUR | 7.53e-08 |
| GWD | STU | 9.99e-01 |
| GWD | TSI | 3.07e-01 |
| GWD | YRI | 1.31e-01 |
| IBS | ITU | 1.67e-01 |
| IBS | JPT | 5.60e-07 |
| IBS | KHV | 1.00e+00 |
| IBS | LWK | 2.40e-02 |
| IBS | MSL | 2.18e-11 |
| IBS | MXL | 2.99e-04 |
| IBS | PEL | 9.99e-01 |
| IBS | PJL | 9.76e-01 |
| IBS | PUR | 9.58e-01 |
| IBS | STU | 5.17e-01 |
| IBS | TSI | 1.35e-10 |
| IBS | YRI | 9.97e-01 |
| ITU | JPT | 8.17e-01 |
| ITU | KHV | 2.78e-01 |
| ITU | LWK | 1.00e+00 |
| ITU | MSL | 4.00e-03 |
| ITU | MXL | 9.01e-10 |
| ITU | PEL | 9.46e-01 |
| ITU | PJL | 2.54e-04 |
| ITU | PUR | 5.67e-04 |
| ITU | STU | 1.00e+00 |
| ITU | TSI | 2.70e-02 |
| ITU | YRI | 9.78e-01 |
| JPT | KHV | 2.13e-07 |
| JPT | LWK | 1.00e+00 |
| JPT | MSL | 7.31e-01 |
| JPT | MXL | 1.38e-13 |
| JPT | PEL | 5.44e-04 |
| JPT | PJL | 1.27e-12 |
| JPT | PUR | 5.28e-11 |
| JPT | STU | 6.12e-01 |
| JPT | TSI | 9.89e-01 |
| JPT | YRI | 1.00e-03 |
| KHV | LWK | 4.60e-02 |
| KHV | MSL | 1.42e-11 |
| KHV | MXL | 4.36e-07 |
| KHV | PEL | 1.00e+00 |
| KHV | PJL | 1.96e-01 |
| KHV | PUR | 3.21e-01 |
| KHV | STU | 7.26e-01 |
| KHV | TSI | 7.17e-11 |
| KHV | YRI | 1.00e+00 |
| LWK | MSL | 5.36e-01 |
| LWK | MXL | 1.15e-09 |
| LWK | PEL | 3.96e-01 |
| LWK | PJL | 7.35e-05 |
| LWK | PUR | 1.05e-04 |
| LWK | STU | 1.00e+00 |
| LWK | TSI | 9.03e-01 |
| LWK | YRI | 4.94e-01 |
| MSL | MXL | 2.54e-14 |
| MSL | PEL | 2.61e-08 |
| MSL | PJL | 2.44e-13 |
| MSL | PUR | 1.67e-13 |
| MSL | STU | 2.00e-03 |
| MSL | TSI | 1.00e+00 |
| MSL | YRI | 6.40e-08 |
| MXL | PEL | 3.63e-07 |
| MXL | PJL | 4.10e-02 |
| MXL | PUR | 3.63e-01 |
| MXL | STU | 6.70e-08 |
| MXL | TSI | 0.00e+00 |
| MXL | YRI | 2.09e-07 |
| PEL | PJL | 8.40e-02 |
| PEL | PUR | 1.25e-01 |
| PEL | STU | 9.98e-01 |
| PEL | TSI | 2.13e-07 |
| PEL | YRI | 1.00e+00 |
| PJL | PUR | 1.00e+00 |
| PJL | STU | 5.00e-03 |
| PJL | TSI | 2.61e-13 |
| PJL | YRI | 6.00e-02 |
| PUR | STU | 7.00e-03 |
| PUR | TSI | 2.52e-13 |
| PUR | YRI | 9.30e-02 |
| STU | TSI | 1.30e-02 |
| STU | YRI | 1.00e+00 |
| TSI | YRI | 5.36e-07 |

**Table S3.** Pairwise comparisons of telomere content variation between superpopulations using Games-Howell post hoc test. Significant differences are highlighted in the table below.

| Comparison | P.adj | Interpretation |
| --- | --- | --- |
| African ancestry vs American ancestry | 5.68e-13 | Significant difference (p <0.05) |
| African Ancestry vs East Asian Ancestry | 1.47e-01 | Non-significant |
| American Ancestry vs East Asian Ancestry | 0.00e+00 | Significant difference (p <0.05) |
| African Ancestry vs European Ancestry | 5.95e-04 | Significant difference (p <0.05) |
| American Ancestry vs European Ancestry | 7.55e-05 | Significant difference (p <0.05) |
| East Asian Ancestry vs European Ancestry | 2.01e-01 | Non-significant |
| African Ancestry vs South Asian Ancestry | 2.02e-01 | Non-significant |
| American Ancestry vs South Asian Ancestry | 0.00e+00 | Significant difference (p <0.05) |
| East Asian Ancestry vs South Asian Ancestry | 1.00e+00 | Non-significant |
| European Ancestry vs South Asian Ancestry | 2.96e-01 | Non-significant |

**Table S4.** Pairwise comparisons of telomere content variation between self- reported males and females within population in the Phase 3 of 1000 Genomes Project data using a Welch’s t-test. The Bonferroni correction was applied to adjust the p-value to correct for multiple testing. Significant differences (uncorrected *p < 0.05*) are highlighted in the table below.

| Population | P_value | P_value_Bonferroni |
| --- | --- | --- |
| ACB | 0.9767105 | 1.000000000 |
| ASW | 0.4372406 | 1.000000000 |
| BEB | 0.2159207 | 1.000000000 |
| CDX | 0.0001872209 | 0.004867744 |
| CEU | 0.3615737 | 1.000000000 |
| CHB | 0.003219108 | 0.083696796 |
| CHS | 0.6003528 | 1.000000000 |
| CLM | 0.8481509 | 1.000000000 |
| ESN | 0.9110541 | 1.000000000 |
| FIN | 0.5646176 | 1.000000000 |
| GBR | 0.219772 | 1.000000000 |
| GIH | 0.09874763 | 1.000000000 |
| GWD | 0.7830224 | 1.000000000 |
| IBS | 0.9664038 | 1.000000000 |
| ITU | 0.5644896 | 1.000000000 |
| JPT | 0.9754833 | 1.000000000 |
| KHV | 0.8349956 | 1.000000000 |
| LWK | 0.5572195 | 1.000000000 |
| MSL | 0.4903071 | 1.000000000 |
| MXL | 0.6635297 | 1.000000000 |
| PEL | 0.93373 | 1.000000000 |
| PJL | 0.7490501 | 1.000000000 |
| PUR | 0.708827 | 1.000000000 |
| STU | 0.928914 | 1.000000000 |
| TSI | 0.4402604 | 1.000000000 |
| YRI | 0.2240262 | 1.000000000 |

